# Metabolite annotation from knowns to unknowns through knowledge-guided multi-layer metabolic network

**DOI:** 10.1101/2022.06.02.494523

**Authors:** Zhiwei Zhou, Mingdu Luo, Haosong Zhang, Yandong Yin, Yuping Cai, Zheng-Jiang Zhu

## Abstract

Liquid chromatography - mass spectrometry (LC-MS) based untargeted metabolomics allows to measure both known and unknown metabolites in the metabolome. However, unknown metabolite annotation is a grand challenge in untargeted metabolomics. Here, we developed an approach, namely, knowledge-guided multi-layer network (KGMN), to enable global metabolite annotation from knowns to unknowns in untargeted metabolomics. The KGMN approach integrated three-layer networks, including knowledge-based metabolic reaction network, knowledge-guided MS/MS similarity network, and global peak correlation network. To demonstrate the principle, we applied KGMN in an *in-vitro* enzymatic reaction system and different biological samples, with ∼100-300 putative unknowns annotated in each data set. Among them, >80% unknown metabolites were validated with *in-silico* MS/MS tools. Finally, we successfully validated 5 unknown metabolites through the repository-mining and the syntheses of chemical standards. Together, the KGMN approach enables efficient unknown annotations, and substantially advances the discovery of recurrent unknown metabolites towards deciphering dark matters in untargeted metabolomics.

## Introduction

The metabolome refers to the complete collection of small molecules in living organisms^1–4^. It includes not just endogenously produced known metabolites from cellular metabolism, but also unknown metabolites generated from microbiota, plants, foods and xenobiotics^3,5,6^. Liquid chromatography - mass spectrometry (LC-MS) based untargeted metabolomics allows to measure thousands of metabolic features from biological samples^7,8^. These metabolic features come from known and unknown metabolites, as well as abiotic MS signals generated during ionization such as adducts, isotopes, neutral losses and other ions generated from in-source fragmentation^9,10^. Metabolite identification remains the central bottleneck in LC-MS based untargeted metabolomics^4,11^. For annotation of known metabolites, the most common approach is to search the exact mass of precursor ion (MS1 *m/z*) and tandem mass spectrum (MS2 spectrum) against the standard spectral libraries^12,13^. Significant efforts were made in the past decades to expand the coverage of spectral libraries. For annotation of unknown metabolites, due to lacking the knowledge of chemical structures, additional experiments or in-silico tools were generally required^5,11^. For example, Tsugawa and colleagues employed the stable-isotope labeling to determine formulas of unknown metabolites through identifying the labeled and non-labeled pair of metabolic peaks^14^. In addition, bioinformatic tools, such as MetFrag^15^, CFM-ID^16^, MS-FINDER^17^ and SIRIUS^18^, have been developed to predict *in-silico* MS/MS or molecular fingerprinting to elucidate unknowns. These tools largely rely on retrieving putative chemical structures from existing structural databases (e.g. HMDB^19^ and PubChem^20^). Therefore, it is not suitable for identifying unknown metabolites not included in the databases. Alternatively, COSMIC was recently developed to annotate unknowns from *in-silico* generated metabolites^21^. In general, these bioinformatic tools focus on annotating metabolites for individual metabolic peak and MS2 spectrum without considering other MS signals and peaks in the same dataset.

Recently, network-based approaches are increasingly adopted for metabolite annotation, such as GNPS^22^, MetDNA^23^, and NetID^24^. GNPS is one of most popular approaches, which linked similar MS/MS spectra in datasets and facilitated to infer structurally similar metabolites without standard MS2 spectra available. Other *in-silico* approaches embedded in GNPS further enhanced the capability to elucidate the metabolite structures, including NAP^25^, MS2LDA^26^, and MolNetEnhancer^27^. Recently, NetID was developed to use the integer linear programming approach to optimize a peak correlation network^24^. NetID improved the accuracy of peak assignments and provided a possible formula transformation between peaks. Such a formula transformation facilitated the discovery of unknowns. Although these methods have proved their effectiveness for the discovery of unknown metabolites, professional mass spectrometrists are still required to manually curate the chemical structures of unknown metabolites, which restricts the annotation efficiency of unknown metabolites with a global scale. As a comparison, MetDNA utilized the metabolic reaction network (MRN) and MS/MS spectral similarity to connect metabolic peaks with a recursive manner, achieving high-coverage and efficient metabolite annotation^23^. However, it cannot support unknown metabolites which are not covered in metabolic reaction network.

In this work, we developed an approach, namely, knowledge-guided multi-layer network (KGMN), to enable global metabolite annotation from knowns to unknowns in untargeted metabolomics (**Figure 1**). The KGMN approach integrated three layers of networks, including knowledge-based metabolic reaction network (KMRN), knowledge-guided MS2 similarity network, and global peak correlation network. We first demonstrated that the multi-layer metabolic network strategy significantly improved the identification accuracy of known metabolites to >95% compared to MetDNA, in which only a single metabolic reaction network was used. Furthermore, we demonstrated the principle of metabolite annotation from knowns to unknowns using KGMN in an *in-vitro* enzymatic reaction system and different biological samples, with ∼100-300 putative unknowns annotated in each data set. Most importantly, more than 80% unknown metabolites were validated with other *in-silico* MS/MS tools. Finally, we successfully discovered 5 unknown metabolites through integrating KGMN and the repository-mining. Altogether, the KGMN approach allows efficient annotations of unknown metabolites and substantially advances the discovery of recurrent unknowns towards deciphering dark matters in untargeted metabolomics.

**Figure 1.**
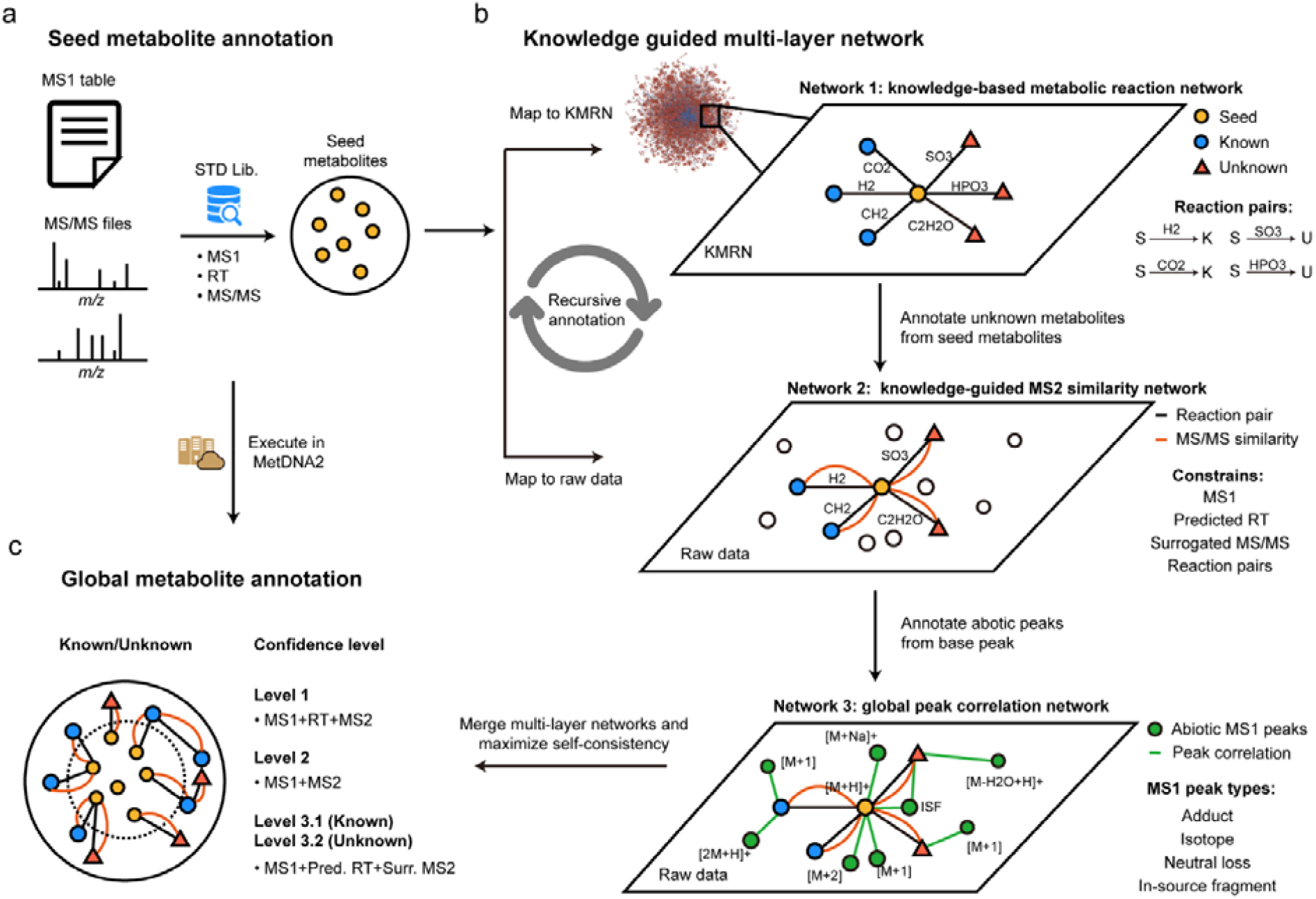
The workflow of knowledge-guided multi-layer network (KGMN) for metabolite annotations in untargeted metabolomics. (**a**) Annotation of seed metabolites by matching with the standard library using MS1, retention time (RT), and MS/MS spectra. (**b**) Knowledge-guided multi-layer network enables global metabolite annotations by propagation from knowns to unknowns. The network 1 is a knowledge-based metabolic reaction network (KMRN), where known or unknown metabolites are linked by known and *in-silico* metabolic reactions. The network 2 is a knowledge-guided MS2 similarity network, where annotations propagate from known to unknown along KMRN. The network 3 is a global peak correlation network to recognize abiotic peaks and improve the accuracy of peak assignment. (**c**) The KGMN based metabolite annotations are executed in an automated and unsupervised manner and reported with definitive confidence levels.

## Results

### The workflow

Knowledge-guided multi-layer network enables global unknown metabolite annotation by propagating annotations from knowns to unknowns along metabolic reaction network (**Figure 1**). The KGMN approach integrated three layers of networks: (1) knowledge-based metabolic reaction network (KMRN), (2) knowledge-guided MS2 similarity network, and (3) global peak correlation network. The seed metabolites were first annotated by matching their properties (MS1, retention time, and MS2 spectra) to metabolite standard libraries (**Figure 1a**), and mapped into metabolic reaction network to retrieve reaction-paired neighbor metabolites (network 1 in **Figure 1b**). This network is a knowledge-based metabolic reaction network, where known or unknown metabolites are linked by either known or in-silico reactions. Specific to unknowns, they are curated via performing *in-silico* enzymatic reactions using known metabolites in KEGG database as substrates (see Methods and **Extended Data Fig. 1**). For example, oxaloacetate (C00036) can be reduced to malate (C00149) with a reductase. Such a reduction reaction could be applied to other structurally similar metabolites and generate possible unknowns with novel structures (**Extended Data Fig. 1a**). These unknown products are linked with their precursors to expand metabolic network from known chemical space to unknown space. In sum, a total of 34,858 unknown metabolites were generated from known metabolites, and further linked together through 52,137 edges and 1,504 biotransformation types in KMRN (**Extended Data Fig. 1b-d**). These unknown metabolites included 405 known-unknowns and 32,980 unknown-unknowns depending on whether they were included by HMDB (version 4.0, released on 2018-12-18).

KMRN provides a known-to-unknown based structural similarity network to guide the construction of the MS2 similarity network from the LC-MS/MS data (network 2 in **Figure 1b**). Specifically, reaction-paired neighbor metabolites (knowns or unknowns) from seeds were putatively annotated from the LC-MS/MS data. Their calculated MS1 *m/z* and predicted RTs were matched to the experimental values in the data file. Meanwhile, surrogated MS2 spectrum from seed metabolite was used for MS/MS spectral match. The matched peaks were annotated as putative neighbor metabolites and linked to seeds in network 2. Thereby, seeds linked to other annotated metabolites with 4 constrains, including MS1 *m/z*, RT, MS/MS similarity, and a metabolic biotransformation (e.g., reduction, +2H; decarboxylation, -CO_2_). Compared to GNPS and other tools which solely use the MS/MS similarity to construct the network, our knowledge-guided MS2 similarity network have explicable structural relationships between two nodes and more succinct network topology (**Supplementary Figure. 1**). In addition, similar to MetDNA, the annotated metabolites could be used as new seeds to annotate more metabolites and extend the network. Such annotation is performed in a recursive manner until there is no new metabolites annotated in LC-MS/MS data.

The third global peak correlation network aims to annotate abiotic peaks in LC-MS data (network 3 in **Figure 1b**). All annotated peaks from MS2 similarity network were considered as base peaks. Next, abiotic peaks derived from each base peak were targeted extracted from the peak list in LC-MS data, including adducts, isotopes, neutral losses, and other in-source fragments (ISF). More specifically, common adducts (e.g. Na^+^, K^+^) and neutral losses (e.g. -H_2_O, -NH_3_) were searched within co-eluted peaks, while in-source fragments were retrieved from the MS2 spectra of the base peaks (See Methods and **Extended Data Fig. 2**). Then, a peak correlation subnetwork is constructed for each annotated metabolite through connecting base peak and abiotic peaks. The subnetwork describes the comprehensive peak profiles of the metabolite during ionization in mass spectrometry measurements. As a result, a global peak correlation network (network 3) is constructed through combining the subnetworks of all base peaks and the knowledge-guided MS2 similarity network. Similar to NetID, the global peak correlation network provides a valuable basis to optimize metabolite annotation from the first two-layer network and improve the accuracy of peak assignment. Peak annotations are compared and scored within and across subnetworks, conflicts are further resolved by maximizing the self-consistency in each subnetwork. The subnetworks with most linked edges were reserved while unsatisfactory and conflict subnetworks and their putative annotations were removed.

Finally, the annotation results from KGMN were outputted with definitive confidence levels according to the MSI guidelines^28^ (**Figure 1c**). The KGMN approach is implemented and freely available in MetDNA2 webserver (http://metdna.zhulab.cn/). The KGMN accepts various data imports from common data processing tools, including XCMS^29^, MS-DIAL^30^, and MZmine2^31^. The step-by-step tutorials are provided in the website.

### KGMN improves peak annotation accuracy

The KGMN approach enables to optimize metabolite annotation and improve the accuracy of peak assignment through global peak correlation network. Here, we demonstrate the principle with an example in NIST urine sample (**Figure 2a**). Metabolic features M285T555 and M153T555 were putatively annotated as xanthosine and 5-ureido-4-imidazole carboxylate, respectively. The base peaks of both metabolites and their related abiotic peaks were extracted to construct subnetworks. Two subnetworks had 14 and 7 recognized peaks, respectively. One conflict peak assignment was observed in these subnetworks. The base feature of M153T555 is assigned as an in-source fragment ion of xanthosine. To resolve the conflict and maximize the self-consistency of peak annotations in two subnetworks, the subnetwork of M153T555 and its annotation of 5-ureido-4-imidazole were removed. As a result, all peak annotations in two subnetworks could be merged and confirmed to be associated with xanthosine. We further validated that M153T555 is an in-source fragment of xanthosine using the chemical standard (**Figure 2b**).

**Figure 2.**
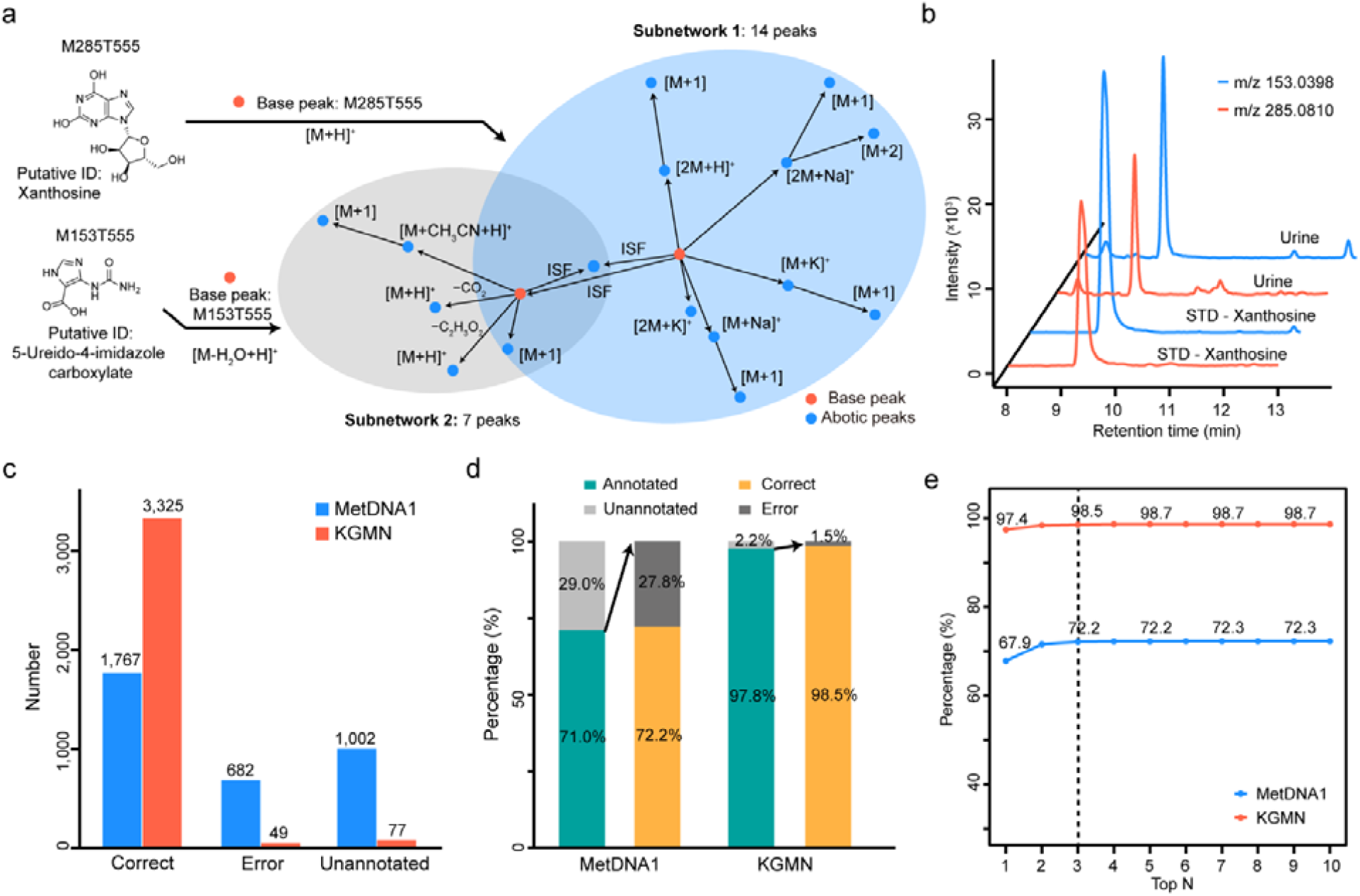
KGMN improves peak annotation accuracy. **(a**) Peak correlation subnetworks for metabolic features of M285T555 and M153T555. The M153T555 was correctly recognized as the in-source fragment of M285T555 by KGMN. (**b**) The validation of M153T555 (m/z 153.0398) as an in-source fragment of M285T555 (m/z 285.0810) using the chemical standard of xanthiosine. (**c-e**) The improved accuracy of known metabolite annotation using MetDNA1 and KGMN: **c**, statistics of annotation numbers; **d**, percentages of annotation coverage and correct/error rates; **e**, correct and error rates among top n annotations. The x-axis represents the top n metabolite candidates for each peak.

Then, we systemically evaluated the improved accuracy of peak annotations with a manually curated data set that contains a total of 242 metabolites, 3,451 metabolic features from 5 different biological samples and 2 ionization polarities (**Supplementary Table 1** and **Supplementary Data 1**). Among these metabolic features, our previously developed MetDNA (denoted as MetDNA1) reported a total of 2,449 annotations, including 1,767 correct (72.2%) and 682 error (27.8%) annotations. The remaining 1,002 peaks were not annotated (**Figure 2c**). As a comparison, with the optimization of global peak correlation network, the KGMN approach significantly increased the correct peak annotations to 3,325 (98.5%) and decreased error annotations to 49 (1.5%; **Figure 2c**). As shown in **Figure 2d**, when only the annotated peaks were considered, the annotation coverage increased from 71.0% to 97.8%, while the correct peak annotations increased from 72.2% to 98.5%. Considering different metabolite annotations for one peak, the correct annotation rates were also consistently improved (**Figure 2e**). Similar results were also obtained for individual data sets in both positive and negative modes (**Supplementary Figure. 2**). These results demonstrated that the KGMN approach effectively extended annotation coverage and increased annotation accuracy.

The most important feature of KGMN is that the peak assignment evaluation and optimization is performed for all peaks in an automated and unsupervised manner, and no manual analyses were required. In addition, we found this approach highly effective to recognize false positive annotations caused by in-source fragments. For example, in-source fragment M112T282 was annotated as the metabolite cytosine in MetDNA1 because of its high MS/MS score (0.9817) (**Extended Data Fig. 3**). This feature was successfully recognized as an in-source fragment feature of the metabolite N4-acetylcytidine by the KGMN approach. More examples were provided in the **Extended Data Fig. 4**. Taken together, the results validated that our KGMN approach provides substantial improvements of peak assignment accuracy, which will facilitate accurate annotations of unknown metabolites in complex biological samples.

### Metabolite annotation from knowns to unknowns

To demonstrate the principle of metabolite annotation from knowns to unknowns in KGMN, we experimentally incubated a mixture of 46 common metabolites (46std_mix) with the human liver S9 fraction for 24 h (**Figure 3a**). The liver S9 fraction contains most phase I and phase II metabolic enzymes, and is widely used to investigate *in-vitro* metabolism. Here, we treated the 46 compounds as known seed metabolites, while their *in-vitro* metabolic products are defined as unknowns. The unknown metabolites in the incubation solution were analyzed by LC-MS/MS. To identify unknowns, we constructed the knowledge-based metabolic reaction network from 46 metabolites, including 531 possible unknown structures and 642 reaction pairs (**Extended Data Fig. 5** and **Supplementary Data 2**). This knowledge-based metabolic reaction network was used to annotate unknown metabolites in LC-MS/MS data. In positive mode, the KGMN approach annotated 36 knowns and 94 unknown peaks, while a total of 745 MS1 peaks associated with known and unknown metabolites were discovered in global peak correlation network (**Figure 3b, Extended Data Fig. 5** and **Supplementary Data 3**). The unknown metabolites were generated from 9 types of biotransformation (**Supplementary Table 2**). Similar results were also obtained in negative mode (**Figure 3c**). It annotated 37 known and 164 unknown peaks, and a total of 932 peaks in global peak correlation network. The unknown metabolites were generated from 15 types of biotransformation (**Supplementary Table 2**).

**Figure 3.**
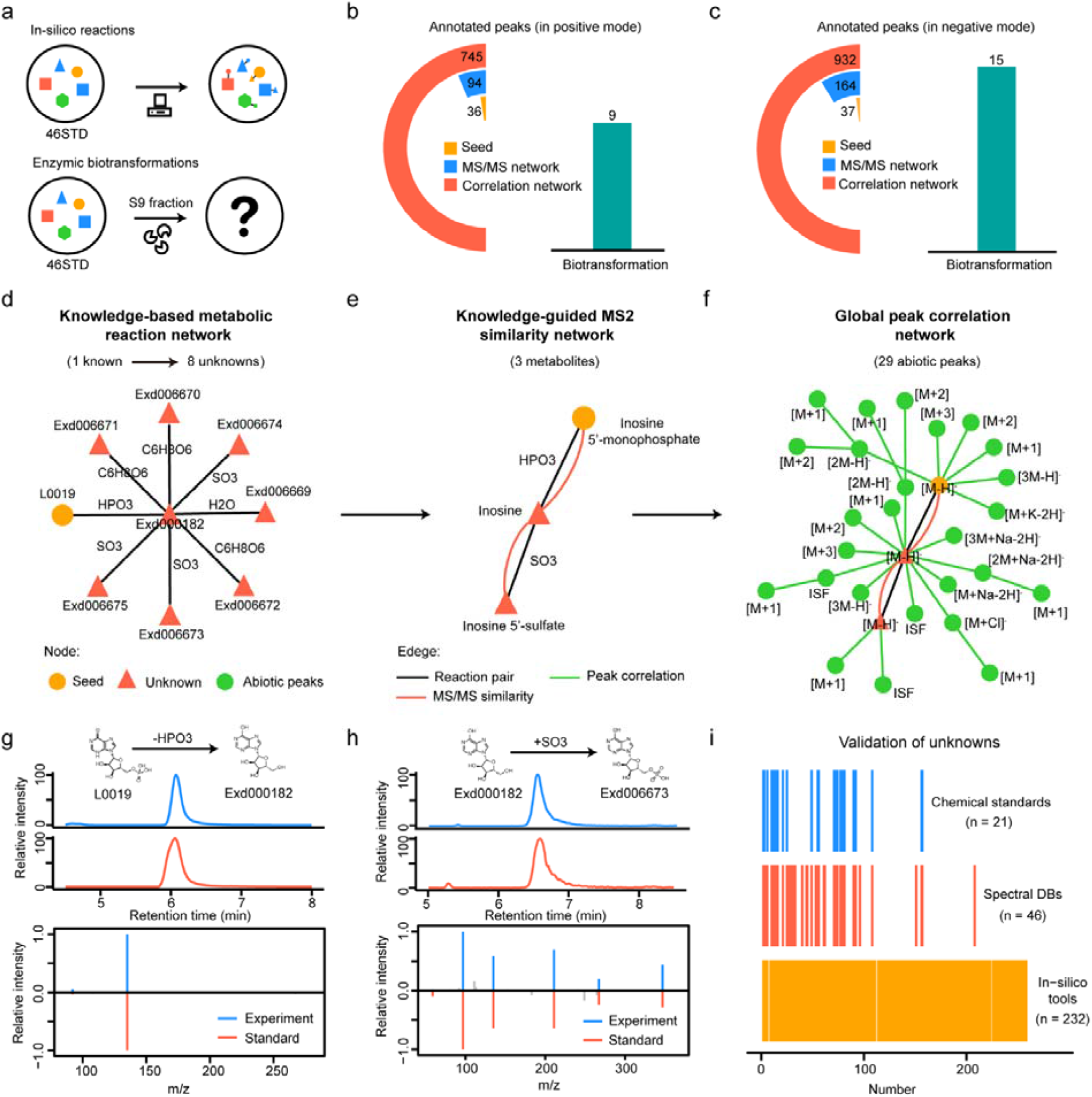
Metabolite annotation from knowns to unknowns. (**a**) The generations of unknown metabolites from 46 metabolite mixtures (46std_mix) with *in-silico* reactions or enzymatic biotransformation via human liver S9 fraction incubation. (**b-c**) The annotated peaks in positive (**b**) and negative modes (**c**); the left cyclic bars represent the annotated peaks in different networks. The right bar represents the involved biotransformation in unknown annotation. (**d**-**f**) The unknown annotations from seed metabolite inosine 5’-monophosphate (IMP, L0019): (**d**) IMP generates 8 unknowns through 4 transformations in knowledge-based metabolic reaction network; (**e**) knowledge-guided MS2 similarity network annotates 2 unknowns from the seed; (**f**) 29 abiotic peaks were annotated from 3 metabolites in global peak correlation network. (**g-h**) Validation of two annotated unknowns: inosine (**f**, labeled as Exd000182) and inosine 5’-sulfate (**g**, labeled as Exd006673). (**i**) The validation of annotated unknowns with different strategies.

We further demonstrated the known-to-unknown annotation with an example of inosine 5’-monophosphate (IMP, denoted as L0019 in **Figure 3d-f**). As one of the seed metabolites, the IMP was identified through matching the standard MS/MS spectral library. Then, the IMP-related-subnetwork was retrieved from knowledge-based metabolic reaction network (**Figure 3d**). Specifically, 8 metabolites were *in-silico* generated from IMP through one- or two-step reactions with 4 different types of biotransformation, including dephosphorylation (-HPO_3_), sulfation (+SO_3_), glucuronidation (+C_6_H_8_O_6_) and hydrolysis (-H_2_O). With the guidance of IMP-reaction network, the knowledge-guided MS2 similarity network was constructed (**Figure 3e**). In the network, 3 metabolites were connected via reaction pair (black line) and MS/MS similarity (orange line), including 1 seed metabolite (IMP) and 2 unknowns (inosine and inosine 5’-sulfate). Furthermore, a peak correlation network was constructed with 29 assigned abiotic peaks from 3 annotated metabolites (**Figure 3f**). In this example, inosine is the product from the dephosphorylation of IMP, and further converted to inosine 5’-sulfate through sulfation. We further confirmed the identification of inosine and inosine 5’-sulfate with the chemical standards (**Figure 3g and 3h**).

Finally, we validated the accuracy of annotated unknown metabolites with multiple strategies, including chemical standards, public spectral libraries, and *in-silico* MS/MS tools (**Figure 3i** and **Supplementary Data 3**). For all 258 annotated unknowns in positive and negative modes (**Figure 3a** and **b**), 25 (10%), 46 (18%), and 232 (90%) were validated by the chemical standards, public spectral libraries, and *in-silico* MS/MS tools, respectively. Some detailed examples were also provided in the **Extended Data Fig. 6**. Taken together, the results proved that the KGMN strategy effectively annotated unknown metabolites from knowns and provided reliable putative structures for unknown peaks in a large scale.

### Global annotation of unknown metabolites

To determine the performance for real biological samples, we applied the KGMN workflow to the untargeted metabolomics data of NIST human urine samples. In positive mode, 173 seed metabolites were first annotated through matching with the standard library (**Figure 4a**). Then, a total of 927 peaks including 634 knowns and 293 unknowns were annotated through knowledge-guided MS/MS similarity network (**Extended Data Fig. 7**). Finally, 3,301 MS1 peaks associated with metabolites were annotated in global peak correlation network (**Extended Data Fig. 8**). The confidence levels of putatively annotated knowns and unknowns through our KGMN approach were assigned as level 3.1 and 3.2, respectively (**Figure 4a**). Similarly, in negative mode, 161 seed metabolites were first annotated while 1,283 peaks including 652 knowns and 631 unknowns were further annotated through knowledge-guided MS/MS similarity network (**Extended Data Fig. 9**). These results demonstrated that the KGMN approach significantly expands the metabolite annotation coverage from seed metabolites in biological samples. In this work, we pay attention to these putative unknown metabolites. Since unknowns are not included in spectra database and lack chemical standards, we employed common *in-silico* MS/MS tools to validate their reliability (**Supplementary Data 4**). Specifically, 206, 95, and 196 unknown peaks in positive mode, and 540, 309, 299 unknown peaks in negative mode were validated by MetFrag^15^, CFM-ID^16^ and MS-FINDER^17^, respectively (**Figure 4b** and **Extended Data Fig. 9**). In sum, 237 (80.9%) and 547 (86.7%) unknown peaks were validated by at least one *in-silico* MS/MS tool in positive and negative modes, respectively. For example, the glycine-conjugated metabolite, 4-hydroxyhippuric acid was validated by *in-silico* MS/MS tools (**Figure 4c**). More validation examples were also provided in **Extended Data Fig. 9**.

**Figure 4.**
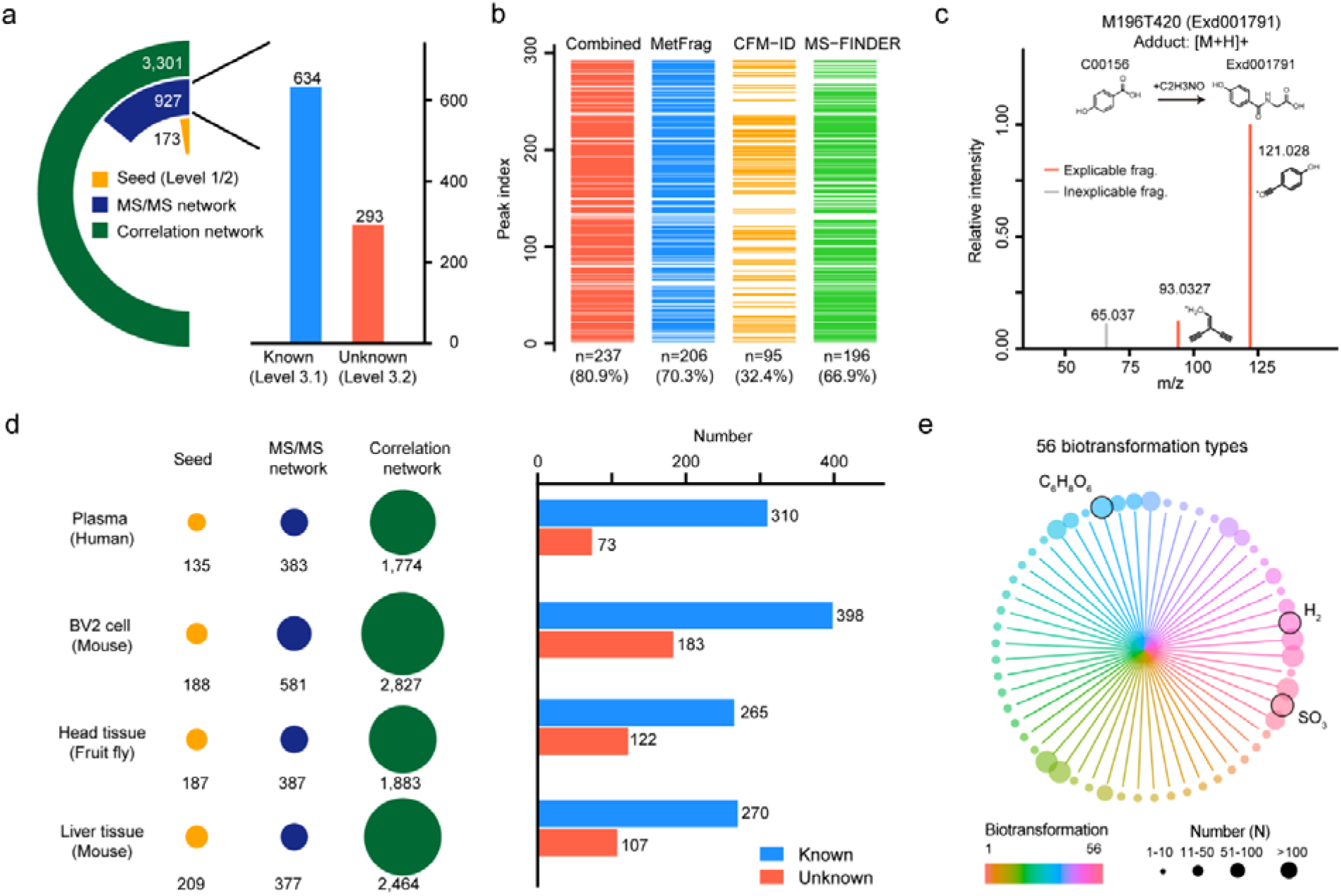
Global annotation of unknown metabolites in biological samples. (**a**) The annotated known and unknown metabolites in NIST human urine samples (positive mode). The left panel is the statistics of annotated peaks in the multi-layer networks, and the right panel is the statistics of annotated known and unknown peaks. (**b**) Validations of annotated unknown metabolites with different *in-silico* MS/MS tools. (**c**) A validation example of the unknown metabolite 4-hydroxyhippuric acid using *in-silico* MS/MS tools. (**d**) Global annotations of metabolites of different biological samples in positive mode. The left panel is the statistics of annotated peaks in the multi-layer network, and the right panel is the statistics of known and unknown metabolites. (**e**) Summary of biotransformation types of annotated unknown metabolites. The color represents different biotransformation types, and the node size represents the frequency number.

Finally, we applied KGMN to different biological samples, including human plasma, BV2 cell, fruit fly head tissue, and mouse liver tissue samples. Consistently, about 100-200 (154±36, Mean ± S.D.), 300-600 (607± 287), and 2000-3000 (2445±758) peaks were annotated in seed annotation, knowledge-guided MS/MS similarity network, and global peak correlation network in positive mode, respectively (**Figure 4d; Supplementary Table 3** and **Supplementary Data 5**). Similar results were also obtained in negative mode (**Extended Data Fig. 9**). On average, ∼100-300 unknown metabolites were annotated in each data sets. These metabolites were generated through 56 types of biotransformation (**Figure 4g** and **Supplementary Table 4**). The most frequent biotransformation types include glucuronidation (C_6_H_8_O_6_), sulfation (SO_3_), and oxidation/reduction (H_2_). Overall, these results demonstrated that the KGMN approach enables global and efficient annotation of unknown metabolites in different biological samples.

### Validation of recurrent unknowns through the repository-mining

With the global annotation of unknown metabolites, it is feasible to evaluate the recurrence of unknowns in public metabolomics data repository. Here, we searched our putatively annotated unknowns in NIST human urine samples against with GNPS/MassIVE database through MASST^32^ (**Figure 5a**). A total of 187 unknowns were recurrent in a total of 351 data sets and 13,640 data files (**Figure 5b** and **Supplementary Data 6**). Specifically, 69, 73, 20 and 25 unknowns were detected in 1 data set, 2-5 data sets, 6-10 datasets, and >10 datasets, respectively. Among them, 76, 25, 13 and 73 unknowns were appeared in 1-10 data files, 11-30 data files, 31-50 data files, and >50 data files, respectively. Furthermore, these datasets originated from 10 different species and 12 different sample types (**Figure 5c**). We observed that recurrent unknown metabolites were enriched on human species, and on plasma/serum and urine, which is consistent with our sample type.

**Figure 5.**
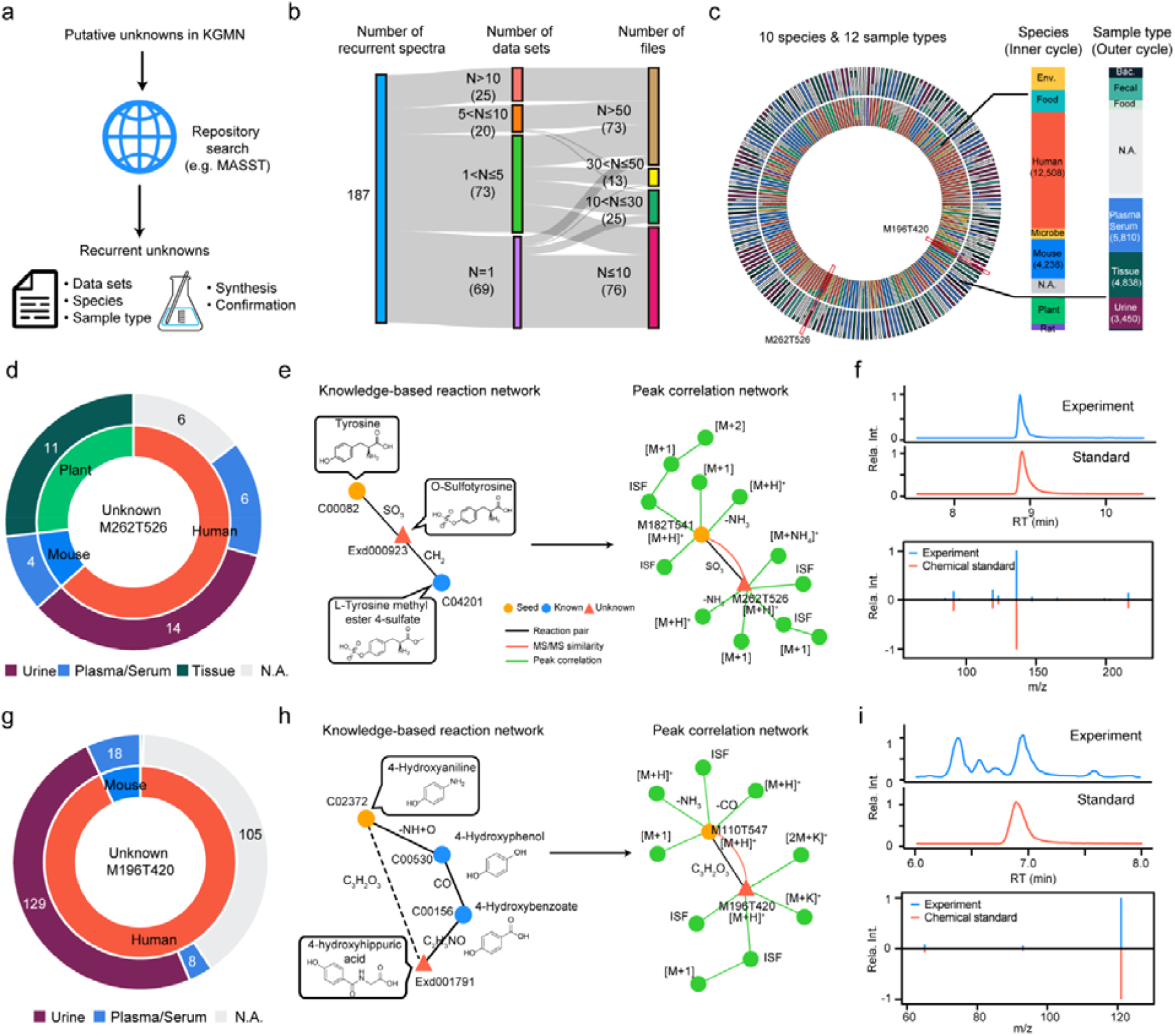
Validation of recurrent unknowns through the repository-mining. (**a**) The workflow for validation of recurrent unknown metabolites using the repository-mining. (**b**) Repository-mining statistics of data sets and data files for 186 recurrent unknown metabolites in human urine samples. (**c**) Repository-mining distributions of 186 recurrent unknown metabolites in species and sample types (left panel). The right panel is the summed distributions of 186 recurrent unknowns in species and sample types. (**d**-**f**) Repository-mining and structural validation of a recurrent unknown metabolite (M262T526): (**d)** the recurrent distributions of species and sample types; the inner and outer pie plots are the distributions in species and sample types, respectively; (**e**) the peak of M262T526 was annotated as O-sulfotyrosine by KGMN; (**f**) structural validation using the synthetic standard by matching retention time and MS/MS spectrum. (**g**-**i**) Repository-mining and structural validation of a recurrent unknown metabolite (M196T420): (**g)** the recurrent distributions of species and sample types; (**e**) the peak of M196T420 was annotated as 4-hydroxyhippuic acid by KGMN; (**f**) structural validation using the synthetic standard by matching retention time and MS/MS spectrum.

We further demonstrated an example for the recurrent unknown peak of M262T526, which was observed in 8 data sets and 45 data files (**Figure 5d**). Through mining their related meta information in GNPS/MassIVE, we found it was reported in multiple species as an unknown, including human (63%), mouse (8%), and plants (27%, e.g. *solanum lycopersicum*). Interestingly, it was only observed in body fluid (plasma, serum, and urine) instead of tissues in mammal. These indicated that this unknown may come from microbiota or xenobiotic resource (e.g. foods). Our KGMN approach putatively annotated this feature as O-sulfotyrosine (**Figure 5e**). In knowledge-based metabolic reaction network, this metabolite can be converted from 2 possible routes, including sulfation (+SO_3_) of tyrosine and demethylation (-CH_2_) of L-tyrosine methyl ester 4-sulfate. In the metabolomics data, tyrosine was firstly annotated in seed annotation, and its annotation further propagated to O-sulfotyrosine with guidance of knowledge-based metabolic reaction network. Finally, O-sulfotyrosine was annotated with 6 related abiotic peaks in its subnetwork through the global peak correlation network. To confirm the unknown annotation, we chemically synthesized O-sulfotyrosine. The synthetic O-sulfotyrosine showed good consistent in both retention time and MS/MS spectrum with the unknown peak in human urine sample (**Figure 5f**). When we are preparing our manuscript, this metabolite is not included in common metabolite databases such as KEGG, HMDB (v4.0), MoNA and GNPS, and therefore can be considered as an unknown-unknown metabolite (**Supplementary Table 5**). O-sulfated metabolites are products of co-metabolism of microbes and their hosts and functional as a class of key regulators for the interaction between microbes and their hosts. This is consistent with the recurrent distributions of this metabolite (**Figure 5d**). Another example of unknown peak M196T420 was only observed human and mouse (**Figure 5g**), which were further annotated as a known-unknown metabolite of 4-hydroxyhippuric acid from the seed metabolite 4-hydroxyanilian in KGMN (**Figure 5h**). Similarly, the annotation was validated using the chemical standard (**Figure 5f**). We also validated another 3 known unknown metabolites in **Extended Fig. 10**. The detailed search history of above known-unknown and unknown-unknown metabolites was provided in **Supplementary Table 5**. Taken together, we demonstrated that the combination of KGMN and the repository-mining facilitated the validation of recurrent unknowns and advanced the understanding of the potential origins and biological functions of newly discovered unknown metabolites.

## Discussion

In this work, we developed the knowledge-guided multi-layer network approach to enable global metabolite annotation from knowns to unknowns in untargeted metabolomics. We demonstrated that the KGMN approach substantially advances the discovery of unknown metabolites towards deciphering dark matters in biological samples. The key advancement of KGMN comes from the proper integration of three metabolic networks to improve the annotation accuracy and enable global unknown annotation. The knowledge-based metabolic reaction network provides a known-to-unknown based structural similarity network to guide the construction of the MS2 similarity network from the LC-MS/MS data. Unknown metabolites are curated via performing *in-silico* enzymatic reactions using known metabolites. Although several publications have reported the use of *in-silico* reactions to annotate unknowns in biological samples, these curated unknowns were simply used as an alternative compound database in these studies^21,33^. As a comparison, KGMN is unique to utilize the knowledge of reaction relationship between reactants and products (*i*.*e*., predicted reaction pair). Importantly, the predicted reaction pairs can further be linked with classic metabolic reaction network (*i*.*e*., KMRN), providing essential routines for annotation from knowns to unknowns. Such knowledge-based reaction network provides straightforward interpretations of unknown annotation and effectively reduced the complexity of MS/MS similarity network generated from untargeted metabolomics data (**Supplementary Figure 1**). Finally, the global peak correlation network is constructed to recognize all possible abiotic peaks, which increases the coverage of recognized peak number by ∼4-5 folds and further strengthens the accuracy of peak assignments. Through integrating the results from above multi-layer networks, our KGMN approach enables the global unknown metabolite annotation in an automated and unsupervised manner.

The accuracy of unknown annotation in KGMN depends on several key factors. First, the characterization of MS/MS spectral similarity has a significant impact on the accuracy of unknown annotation. Specifically, although ∼60% reaction-paired metabolites (either known or predicted pair) shows MS/MS similarity larger than 0.5 (dot product score), the remaining ∼40% reaction pairs have low MS/MS similarity score even they have high structural similarity (**Supplementary Figure 3**). Recently, the similar conclusion was also reported by the Hofft group^34^. We believe the addition of newly developed scoring approaches for MS/MS similarity, like CSS score^35^, Spec2Vec^34^, spectral entropy score^36^, would further enhance the performance of KGMN. Second, we need to be aware of the challenge of discriminating high structurally similar isomers of unknown metabolites. Specific to KGMN, one known metabolite may generate several possible unknown isomers through one biotransformation when performing *in-silico* reaction. For example, isocitrate have 4 reactive function groups for glucuronidation, whose product isomers cannot be distinguished effectively in KGMN. To address these challenges, incorporating more orthogonal properties, such as collision cross-section (CCS)^37^, would be valuable in the future. Third, we think the accuracy of chemical structures of unknowns is largely dependent on the *in-silico* biotransformation algorithms. With continuing innovations of *in-silico* reaction tools (e.g. ATLAS^38^, CyProduct^39^, and MINE2^40^), the structural reliability of predicted unknown metabolites will be improved and thereby benefit the performance of our KGMN approach.

Since unknown metabolites are not included in spectral database and lack chemical standards, the validation of unknown annotation remains a grand challenge. In this work, we first demonstrated the principle of metabolite annotation from knowns to unknowns in an *in-vitro* enzymatic reaction system, and validated the principle with metabolite standards. For unknown annotation in biological samples, we also employed common *in-silico* bioinformatics tools to validate the structural reliability in a large scale. Even so, it is noteworthy that the confidence levels of all annotated metabolites in KGMN belong to level 3 according to the definition of MSI^28^. The metabolite identification still needs to be validated using the synthetic chemical standards. In our work, we demonstrated that the combination of KGMN and the repository-mining facilitated the validation of recurrent unknowns in the repository level. We utilized such an approach to discover 5 recurrent unknown metabolites, and further confirmed their annotations with synthetic chemical standards. With the accumulation of open-source data sets in metabolomics repository, we believe our KGMN approach will pave a new path to validate more unknown structures through the repository-mining.

## Methods

### Chemicals

Pooled human liver S9 fraction (H0610.S9) and NADPH regenerating system (K5100-5) were purchased from Sekisui Xenotech (Kansas City, KS, USA). The cofactors adenosine 3′-phosphate 5′-phosphosulfate lithium salt hydrate (PAPS), acetyl-coenzyme A (acetyl-CoA), uridine diphosphate glucuronic acid (UDPGA) were purchased from Sigma-Aldrich (St. Louis, MO, USA). The glutathione (GSH) was purchased from J&K (Shanghai, China). The NIST urine (SRM 3667) and NIST plasma (SRM 1950) sample were purchased from Ango Biotechnology Co. (Shanghai, China). LC–MS grade methanol (MeOH) and water (H_2_O) were purchased from Honeywell (Muskegon, MI, USA). LC–MS grade acetonitrile (ACN) was purchased from Merck (Darmstadt, Germany). Ammonium hydroxide (NH_4_OH) and ammonium acetate (NH_4_OAc) were purchased from Sigma (St. Louis, MO, USA). Other chemical standards were purchased from Sigma-Aldrich (St. Louis, MO), J&K (Shanghai, China), and TopScience (Shanghai, China).

### Standard MS/MS and RT libraries

In-house MS/MS and RT libraries from chemical standards were used for seed metabolite annotation in KGMN. It supports different types of high-resolution mass spectrometers from various vendors (including Sciex, Agilent, Waters, Bruker and ThermoFisher). The curation of MS/MS spectral library curation followed the previous publication^41,42^. Briefly, a total of 868 metabolites and 611 metabolites were acquired with 14 collision energies from Sciex TripleTOF 5600/6600 and ThermoFisher Exploris 480, respectively. Their retention time were also acquired under Waters BEH Amide column (HILIC) and Phenomenex C18 column (reverse phase). The LC details are provided in LC-MS/MS part.

### Knowledge-based metabolic reaction network

Knowledge-based metabolic reaction network (KMRN) is a network containing known and unknown metabolites (nodes), and their reaction relationship (edges) from known reactions or *in-silico* reactions. The known metabolites and their metabolic reactions were directly downloaded from KEGG reaction pair database (KEGG RCLASS) ^43^ on 7^th^ March, 2017. It contains 6,397 known metabolites and 8,129 known reaction pairs, which was described in our previous MetDNA publication^23^. Unknown metabolites were curated from *in-silico* enzymatic reactions with 6,397 known KEGG metabolites (**Extended Data Fig. 1**). The unknown metabolite is defined as the *in-silico* curated metabolites not included in KEGG database. The BioTransformer^44^ (version 1.0.8) was used for *in-silico* enzymatic reactions, and “EC-based transformation” was used for 2-step reactions. All curated metabolites were merged with the first layer of InChIKey (14 characters) to remove the stereoisomers. The chemical elements of unknown metabolites were restricted within ‘CHONPS’. As a result, a total of 50,471 unknown metabolites were curated via 193 chemical reactions and 114 enzymes. These curated unknown metabolites had the higher natural product likeness than those in PubChem (**Supplementary Figure 3**). The unknown metabolites were further paired with its reactant in an *in-silico* enzymatic reaction, and Tanimoto structural similarity between reaction-paired metabolites was calculated. The reaction pairs with Tanimoto structural similarity larger than 0.7 were reserved. The unpaired metabolites are discarded. The Tanimoto structural similarity was calculated based on PubChem molecular fingerprinting via the R package rcdk (version 3.4.7.1). Finally, both known and unknown reaction pairs were integrated to curate the knowledge-based metabolic reaction network, including 41,336 nodes (6,478 knowns, and 34,858 unknows) and 52,137 edges. Comparing to original network, it increased 34,939 nodes (i.e. metabolites) with in-silico reaction. Through searching against HMDB version 4.0 (released on 2018-12-18), these nodes can be classified as 81 known-known (compound is included in KEGG database), 405 known-unknowns (compound is not included in KEGG but included in HMDB), and 32,980 unknown-unknown (compound is not included in KEGG and HMDB). Consist with MetDNA, reaction-paired knowns and unknowns have higher MS/MS similarity than non-reaction pairs (**Supplementary Fig. 3**).

### Knowledge-guided MS/MS similarity network and annotation propagation

The procedures to curate knowledge-guided MS/MS similarity network followed our previous MetDNA publication with some modifications. Briefly, seed metabolites were first annotated using the standard MS/MS and RT libraries. The match tolerances were set as: MS1 match, 15 ppm; RT match, 20 s; MS2 match, 0.8 (dot product score). The adducts of protonation and deprotonation were used in seed annotation in positive and negative modes, respectively. The seed metabolites were then mapped to KMRN to guide the construction of MS/MS similarity network with 4 constrains, including MS1 *m/z*, predicted RT, MS/MS similarity, and a metabolic biotransformation. The random forest model is used for RT prediction, and its parameter is optimized via R package “caret” (version 6.0-90). Then, seed metabolite-paired knowns/unknowns were retrieved from KMRN, and their calculated MS1 *m/z* and predicted RTs were matched with experimental values in LC-MS/MS data. Match tolerances of MS1 *m/z* and predicted RT were set as 15 ppm and 30%. The qualified peaks were further matched their MS/MS spectra against the surrogated MS2 spectra from seed metabolites. The qualified peaks with dot product score larger than 0.5 or matched fragments more than 4 were linked to the seed metabolites, and their putative structures were assigned from the seeds. Such annotation was propagated in a recursive manner, where newly annotated metabolites were also used as seeds to annotate their neighbor metabolites in LC-MS/MS data. The annotation was terminated until no new metabolites were annotated. A total of 11 and 8 common adducts are considered in annotation propagation in positive and negative modes, respectively (**Supplementary Data 7**).

### Global peak correlation network

Global peak correlation network is used to recognize all possible abiotic peaks in LC-MS data, and further to improve the structural assignment (**Extended Data Fig. 2**). All putatively annotated peaks from knowledge-guided MS/MS network were selected as base peaks, and their co-eluted peaks were extracted from the feature table within ±3 s RT window (composed as a peak group). The recognition of abiotic peaks was performed within each peak group to build a subnetwork, including isotopes, adducts, neutral losses, and in-source fragments (ISF). The detailed procedures are described as follows.

#### Isotope peaks

The recognition of isotope peak includes the evaluations of mass deviation and intensity ratio. The pairwise *m/z* distance matrix was first calculated for deviation check. The theoretical m/z of isotopes were calculated as:

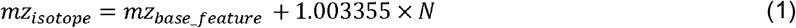

where mz_isotope_ and mz_base_feature_ are *m/z* values for isotopes and base peaks. The N represents considered number of isotopes with a set of values from 1 to 3 (i.e. [M] to [M+3]). The tolerance for mass deviation was set as 25 ppm. The deviation of isotope ratio was calculated as:

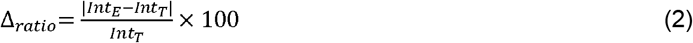

Where Int_E_ and Int_T_ are the experimental and theoretical relative intensities^14^, respectively. The maximum deviation of isotope ratio (^△^_ratio_) was 500% by default.

#### Adduct and neutral loss peaks

A total of 28 types of adducts and 57 types of neutral losses were considered (**Supplementary Data 7**). The adduct and neutral loss features are recognized based on mass deviation and feature abundance correlation among samples. The theoretical *m/z* values of adducts and neutral losses were calculated and matched within each peak group. The tolerance of mass deviation was set as 25 ppm. The feature abundance correlation among samples are calculated between the recognized feature and the base feature, where feature pairs with Pearson correlation coefficient larger than 0.3 are reserved by default. The isotopes of the adduct and neutral loss peaks were also identified using the same approach in “*isotope peaks”*.

#### In-source fragment peaks

The in-source fragment was retrieved from the MS/MS spectrum of base feature and co-eluted MS1 features. Top 5 intense fragments in the MS/MS spectrum of the base peak were considered as possible in-source fragments, and matched with the features in one feature group. The *m/z* tolerance was set as 25 ppm. The isotopes of in-source fragment features were also recognized following the same approach in “*isotope peaks”*.

As a result, a peak correlation subnetwork of one base peak was constructed through connecting base peak and different abiotic peaks. For all base peaks from knowledge-guided MS2 similarity network, and a global peak correlation network was further constructed through combining all subnetworks. Finally, peak recognitions among different subnetworks were compared and optimized to remove the annotation conflicts across the global peak correlation network. Specifically, it contains 3 steps: (1) *check of empirical rules*. The subnetworks are removed if it dissatisfies the empirical rules of relationships between base peaks and its abiotic peaks. The empirical rules are listed in **Supplementary Data 7**. For example, a base peak with the [M+2Na-H]^+^ adduct type requires the presence of an abiotic peak [M+Na]^+^ in its subnetwork; (2) *removal of conflict peaks*. The base peak with different adduct or neural loss annotations is checked, where the subnetwork with larger size is reserved. For example, the base peak have both of [M+H]^+^ and [M-H_2_O+H]^+^ annotations with 5 and 3 abiotic peaks in each subnetwork, respectively. The subnetwork in [M+H]^+^ type is reserved; (3) *consolidation of redundant abiotic peaks*. The multiple abiotic peaks from same metabolite were consolidated to the subnetwork with the maximum size.

### Annotation confidence and reporting

Annotated structures are assigned with different confidence levels according to the definition by Metabolomics Standards Initiative (MSI)^28^. The confidence levels were defined as follows: level 1: metabolites annotated using in-house metabolite standards with three orthogonal properties (*i*.*e*., MS1+RT+MS/MS); level 2: metabolites annotated using two orthogonal properties from the standard MS/MS libraries without RT available (*i*.*e*., MS1 + MS/MS); level 3.1: known KEGG metabolites annotated with MS1, predicted RT, and surrogate MS/MS spectra (*i*.*e*., MS1 + Pred. RT + Surro. MS/MS); level 3.2: unknown structures annotated with MS1, predicted RT, and surrogate MS/MS spectra (*i*.*e*., MS1 + Pred. RT + Surro. MS/MS). For each feature, all candidates were ranked with a total score (S_total_), which was calculated as Eq. 3:

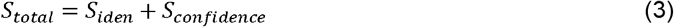

where S_iden_ and S_confidence_ represent the identification score and confidence score, respectively.

The identification score was calculated as Eq. 4:

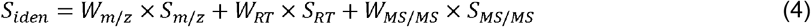

where S_m/z_, S_RT_, and S_MS/MS_ are *m/z* match, RT match and MS/MS match scores, respectively. These scores are calculated using the method as MetDNA. The W_m/z_, W_RT_, W_MS/MS_ are weights for the *m/z* match, RT match and MS/MS match scores, and set as 0.25, 0.25, and 0.5, respectively.

The confidence score was calculated as follows Eq. 5:

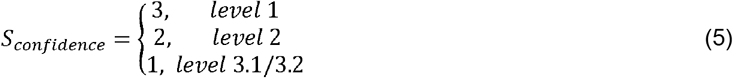

For each feature, annotation candidates with the highest confidence level were reported. If multiple annotations with the same confidence level, top 10 ranked candidates using the total score were kept.

### *In-vitro* metabolism experiment with human liver S9 fraction

We experimentally incubated a mixture of 46 common metabolites (46std_mix) with the human liver S9 fractions for 24 h. The 46std_mix solution was prepared using the concentrations provided in **Supplementary Data 2**, and stored at -80 °C before incubation. The incubation followed the previously reported protocol^45^ with minor modification. Before the experiment, 50 μL of pooled human liver S9 fraction solution (H0610.S9) was diluted into 500 μL using water. The NADPH regenerating system was reconstituted with the addition of 3.5 mL of water to make a final volume of 5 mL. The 4x cofactor stock was freshly prepared with the following composition: 10 mM UDPGA, 2 mM GSH, 2 mg/ml PAPS, 0.1 mM acetyl-CoA, and NADPH regenerating system (1 mM NADP, 5 mM glucose-6-phosphate, 1 unit glucose-6-phosphate dehydrogenase). The incubation was performed in the 1.5 mL of Eppendorf centrifuge tube. In each tube, 30 μL of S9 fraction, 30 μL of Tris buffer (0.2 M; pH 7.5; 2 mM MgCl_2_) and 30 μL of 46std_mix solution were first pooled. To start the reaction, 30 μL of 4× cofactor stock was added, and the incubation was carried out at 30 °C for 24 h. 360 μL of MeOH:ACN (1:1, v:v) were added to terminate the reaction and extract the metabolites. For metabolite extraction, the sample was incubated at -20 °C for 1 hour to facilitate protein precipitation. After the incubation, samples were centrifuged at 13000 rpm and 4 °C for 15 min. The supernatant was taken out and evaporated to dryness at 4 °C. The samples were reconstituted with 120 μL of ACN/H_2_O (v:v, 1:1) and vortexed for 30 s and sonicated for 10 min at 4 °C water bath. Finally, the samples were centrifuged for 15 min at 17,000 xg and 4 °C. The supernatant was taken into the sample vial for LC-MS experiment. Finally, the annotated known and unknown metabolites from incubated 46mix_std samples were validated using multiple strategies. For known metabolite annotations, chemical standards (MS1+RT+MS/MS, level 1) and public spectral database (MS1+ MS/MS, level 2) were used. Public spectral libraries included NIST17, SonnenburgLabLib, Metlin, MassBank, Fiehn HILIC library and GNPS library. For validation of unknown metabolites, 3 different in-silico MS/MS tools were used, including MetFrag (version 2.4.5-CL), CFM-ID (version 2.4) and MS-FINDER (version 3.24).

### Preparation of biological samples

The biological samples were extracted followed our published protocols^46^. In brief, NIST urine samples were thawed at 4 °C on ice. Then 150 μL of urine samples were taken and transferred into a centrifuge tube, and 600 μL of MeOH were added to extract the sample. After vortexed for 30 seconds and sonicated for 10 minutes at 4 °C in water bath, the samples were incubated for 1 hour at -20 °C to facilitate protein precipitation. After the incubation, the samples were further centrifuged for 15 minutes at 17,000 xg and 4 °C. The supernatant was collected and evaporated to dryness at 4 °C. The dry extracts were then reconstituted into 150 μL of ACN:H_2_O (1:1, v/v), followed by sonication at 4 °C for 10 min, and centrifuged at 17,000 xg and 4 °C for 5 min to remove the insoluble debris before LC–MS/MS analysis. For NIST plasma, 100 μL of NIST plasma was extracted using 400 μL of solvent mixture of MeOH:ACN (1:1, v/v) in the centrifuge tube, and then the mixture was vortexed for 30 s and sonicated for 10 min at 4 °C water bath. The rest of the procedure was the same as described for NIST urine sample. For BV2 cells, it was plated in 6-cm dishes at 2000,000 cells/dish, and cultured in DMEM medium containing FBS (10%) and penicillin/streptomycin (1%). The culture medium was quickly removed, and the cells were washed with the cold PBS twice. The cell dishes were placed on dry ice and the metabolite extraction solution (ACN/MeOH/H_2_O=2/2/1, v/v/v, 1 mL) was added to the dishes to quench the metabolism. The extraction solution was pre-cooled at -80 °C for 1 h prior to the extraction. The plates were then incubated at -80 °C for at least 40 min. The cell contents were scraped and transferred to a 1.5 mL Eppendorf tube. The samples were vortexed for 1 min, and centrifuged for 10 min at 17,000 xg and 4°C to precipitate the insoluble materials. The rest of the procedure was the same as described for NIST urine sample.

### LC-MS/MS

LC-MS analysis was performed using a UHPLC system (1290 series; Agilent Technologies, USA) coupled to a quadruple time-of-flight mass spectrometer (TripleTOF 6600, SCIEX). A Waters ACQUITY UPLC BEH Amide column (particle size, 1.7 μm; 100 mm (length) × 2.1 mm (i.d.)) was used for the LC separation and the column temperature was kept at 25 °C. Mobile phase A was 25 mM ammonium hydroxide (NH_4_OH) and 25 mM ammonium acetate (NH_4_OAc) in water, and B was ACN for both the positive and negative modes. The flow rate was 0.3 mL/min and the gradient were set as follows: 0–1 min: 95% B, 1–14 min: 95% B to 65% B, 14–16 min: 65% B to 40% B, 16–18 min: 40% B, 18–18.1 min: 40% B to 95% B, and 18.1–23 min: 95% B. The injection volume was 2 μL. The data acquisition was operated using the information-dependent acquisition (IDA) mode. The source parameters were set as follows: ion source gas 1 (GAS1), 60 psi; ion source gas 2 (GAS2), 60 psi; curtain gas (CUR), 30psi; temperature (TEM), 600° C; declustering potential (DP), 60 V, or −60 V in positive or negative modes, respectively; and ion spray voltage floating (ISVF), 5500 or −4000 V in positive or negative modes, respectively. The TOF MS scan parameters were set as follows: mass range, 60–1200 Da; accumulation time, 200 ms; and dynamic background subtract, on. The product ion scan parameters were set as follows: mass range, 25–1200 Da; accumulation time, 50 ms; collision energy, 30 or −30 V in positive or negative modes, respectively; collision energy spread, 0; resolution, UNIT; charge state, 1 to 1; intensity, 100 cps; exclude isotopes within 4 Da; mass tolerance, 10 ppm; maximum number of candidate ions to monitor per cycle, 6; and exclude former target ions, for 4 s after two occurrences.

### Repository-mining and validation of recurrent unknowns

Recurrent unknowns were obtained through searching MS/MS spectra of putative unknown metabolites against GNPS/MassIV repository-mining via MASST^32^ (http://gnps.ucsd.edu). The data was filtered using the default parameters in GNPS. The mass tolerance for precursor ion and fragment ion was set as 0.01 Da. The library spectra were filtered in the same manner as the input data. All matches between input spectra and library spectra were required to have a score above 0.7 and at least 2 matched peaks. The labels of organs and species are manually added according to the description of projects and their meta information (**Supplementary Data 6**).

Recurrent unknown metabolites are validated with chemical standards through chemical or enzymatic syntheses. *O-sulfotyrosine*: The O-sulfotyrosine was synthesized by MuJin Biotech Inc, Shanghai, China. The chemical structure was confirmed by nuclear magnetic resonance spectroscopy (^1^H NMR, 400 MHz, Methanol-d4), δ(ppm)= 7.26 (s, 4H), 3.78 – 3.49 (m, 1H), 3.03 (t, J = 5.5 Hz, 2H). *4-hydroxyhippuic acid was* synthesized and purchased from Sunway (Shanghai, China). The chemical structure was confirmed by nuclear magnetic resonance spectroscopy (^1^H NMR, 400 MHz, DMSO-d6), δ(ppm) = 12.53 (s, 1H), 10.03 (s, 1H), 8.56 (t, J = 5.9 Hz, 1H), 7.73 (d, J = 8.0 Hz, 2H), 6.81 (d, J = 8.0 Hz, 2H), 3.87 (d, J = 5.8 Hz, 2H). *3-hydroxyhippuic acid* was synthesized and purchased from Macklin (Shanghai, China). The chemical structure was confirmed by nuclear magnetic resonance spectroscopy (^1^H NMR, 400 MHz, DMSO-d6), δ(ppm) = 12.56 (s, 1H), 9.69 (s, 1H), 8.71 (t, J = 5.9 Hz, 1H), 7.41 – 7.09 (m, 3H), 6.92 (dt, J = 6.9, 2.3 Hz, 1H), 3.88 (d, J = 5.8 Hz, 2H). *Protocatechuic acid-3-O-sulfate* and *3-hydroxybenzoic acid 3-O-sulphate* were in-house synthesized using enzymatic reaction (S9 fraction incubation) of their precursors protocatechuic acid and 3-hydroxybenzoate, respectively.

### Validations of unknowns using *in-silico* MS/MS tools

For validation of unknown metabolites in biological samples, 3 different *in-silico* MS/MS tools were used, including MetFrag (version 2.4.5-CL), CFM-ID (version 2.4) and MS-FINDER (version 3.24). Format of imported data and parameters were adjusted according requirements of each tool. The detail parameters were kept the same as our previous publication^37^.

## Data availability

All the metabolomics datasets described in our study can be downloaded from the MetDNA2 website [http://metdna.zhulab.cn/]. The raw data files of NIST human urine, NIST human plasma, and BV2 cell can be accessed at the National Omics Data Encyclopedia [Project ID: OEP003157]. The raw data of in-vitro metabolism can be accessed at National Omics Data Encyclopedia [Project ID: OEP003284]. The raw data of Fruit fly head (Project ID: MTBLS612, MTBLS615) and mouse liver can be accessed at MetaboLights (Project ID: MTBLS601, MTBLS606).

## Code availability

KGMN was mainly developed using R, and is executed in MetDNA2. The source code of MetDNA2 was provided in GitHub [https://github.com/ZhuMetLab/MetDNA2]. The completed functions are provided in the MetDNA2 webserver [http://metdna.zhulab.cn/] via a free registration. The detailed tutorial was also provided in the MetDNA2 webserver.

## Acknowledgements

The work is financially supported by National Natural Science Foundation of China (22022411, 92057114 and 31971356), National Key R&D Program of China (2018YFA0800902), Science and Technology Commission of Shanghai Municipality (21JC1405902), and Shanghai Municipal Science and Technology Major Project (2019SHZDZX02).

## Author contributions

Z.J.Z., Z.Z. and M.L. conceived the idea and designed the algorithm and software. Z.Z. developed the KGMN workflow and MetDNA2 package. M.L. performed the sample preparation, data acquisition and data processing. M.L. and Z.Z. contributed to the tutorial of MetDNA2 webserver. H.Z. contributed to the MASST search and labeling. M.L. and Z.Z. performed the data analysis. Y.Y. contributed to the deploy of MetDNA2 webserver. Y. C. contributed to the preparing of manuscript. Z.Z. and M.L. tested and debugged the program and webserver. Z.J.Z. and Z.Z. wrote the manuscript. Z.J.Z. supervised the project.

## Competing interests

The authors declare no competing interests.

**Extended Data Figure 1.**
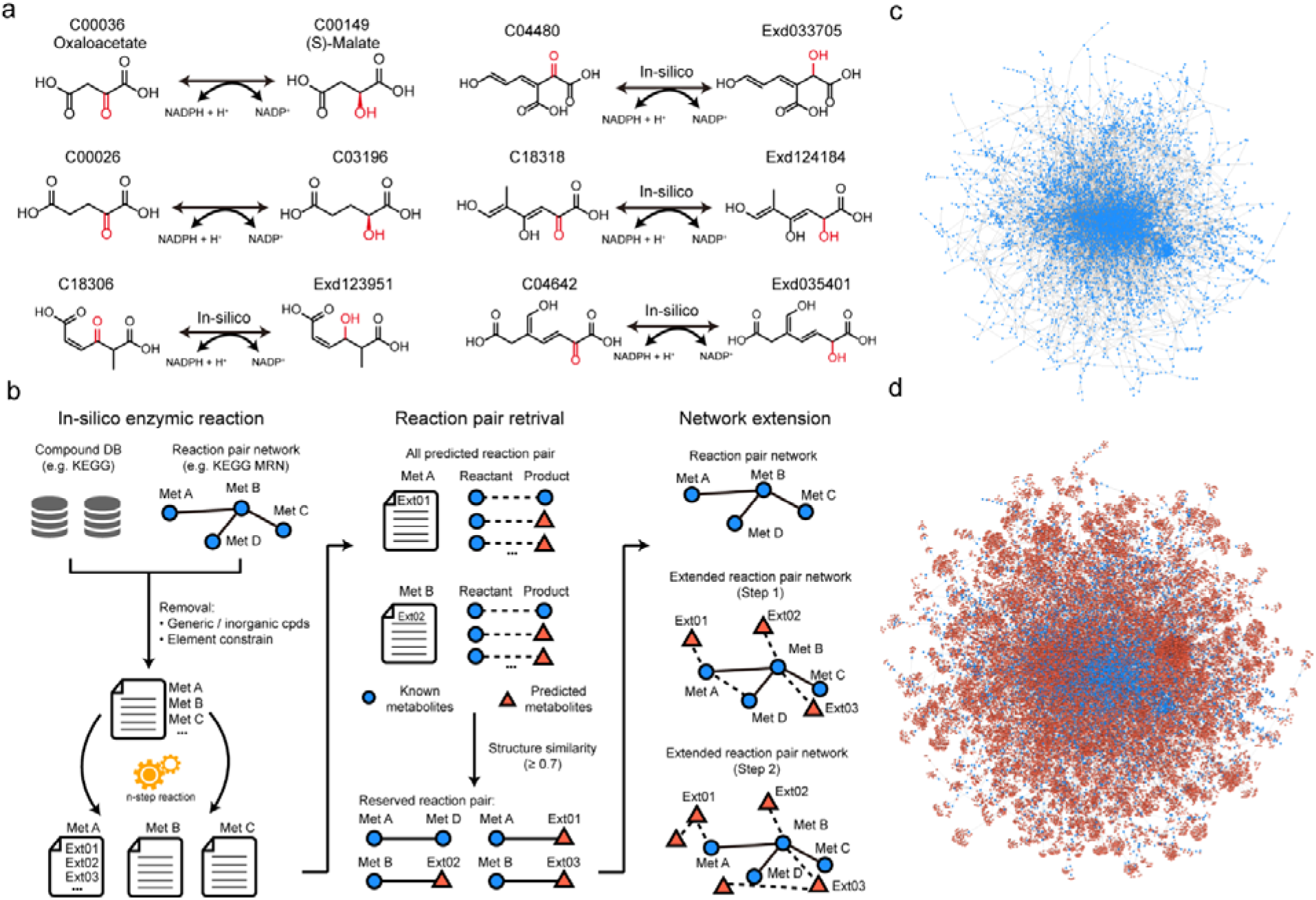
Curation of knowledge-based metabolic reaction network (KMRN) with *in-silico* enzymatic reactions. (**a**) Examples for the curation of unknown metabolites through *in-silico* enzymatic reaction; (**b**) The workflow to curate the knowledge-based metabolic reaction network with *in-silico* enzymatic reactions. The known metabolites and reaction pairs were downloaded from the KEGG database, while the unknown metabolites were curated through *in-silico* enzymatic reactions. The reactant and product were paired and filtered with structural similarity. The knowledge-based metabolic reaction network was linked to the known metabolic reaction network. (**c**-**d**) Knowledge-based metabolic reaction networks: (**c**) known metabolites are connected through known reactions (6,397 nodes and 8,129 edges); (**d**) known and unknown metabolites are connected with known or *in-silico* reactions (41,336 nodes and 52,137 edges). The largest subnetwork is showed here.

**Extended Data Figure 2.**
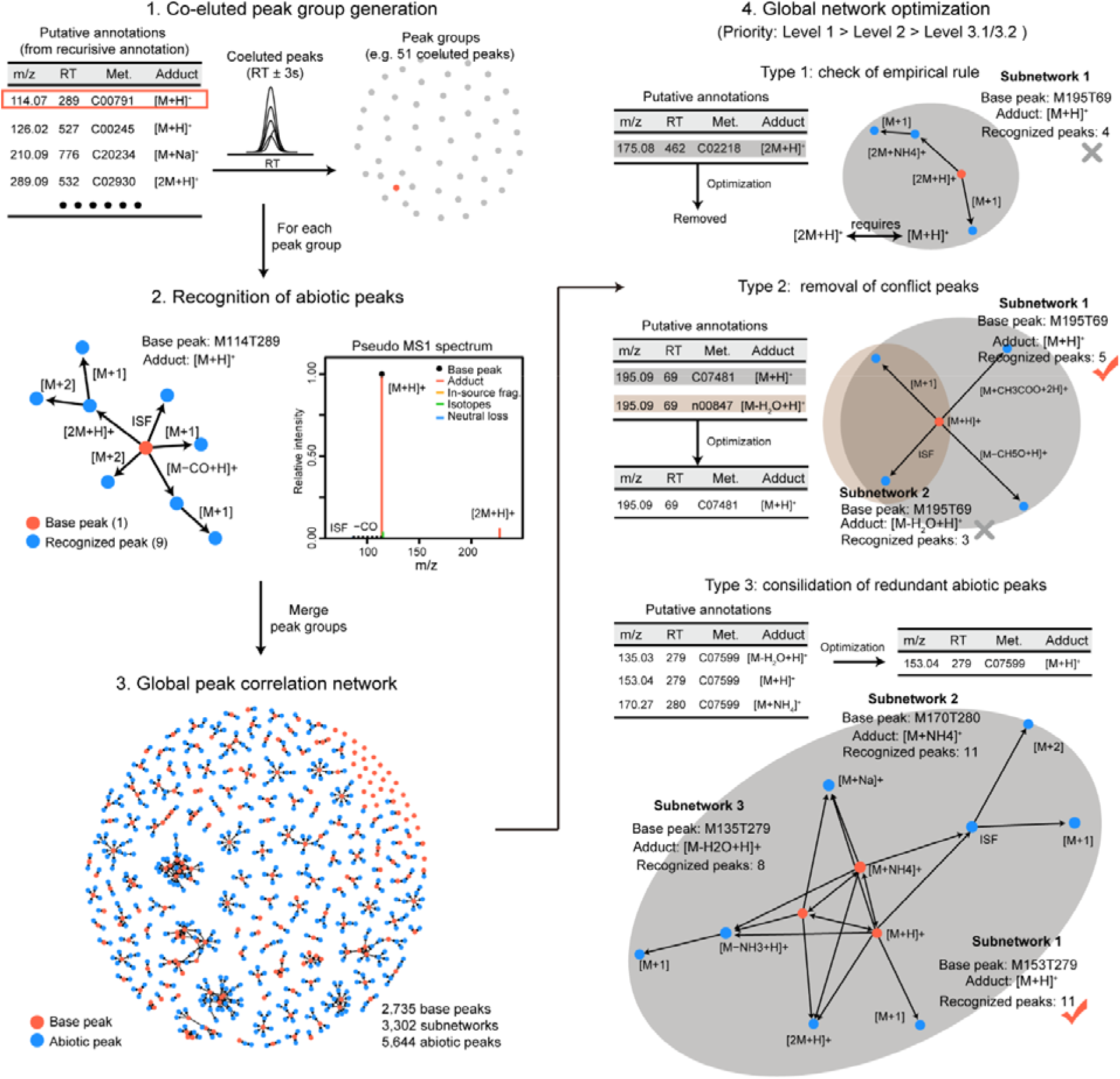
The construction and optimization of global peak annotation network. **Step 1**: co-eluted peaks are extracted as one peak group according to the putative metabolite annotations in knowledge-guided MS/MS similarity network; **Step 2**: recognition of abiotic peaks to build the subnetwork, including adducts, isotopes, in-source fragments and neutral losses; **Step 3**: all recognized subnetworks are merged as a global peak correlation network; **Step 4**: global optimization and conflict resolving to improve the peak annotation accuracy. Three types of conflict annotations are checked and resolved, including empirical rule, removal of conflict peaks and annotations, and consolidation of redundant abiotic peaks.

**Extended Data Figure 3.**
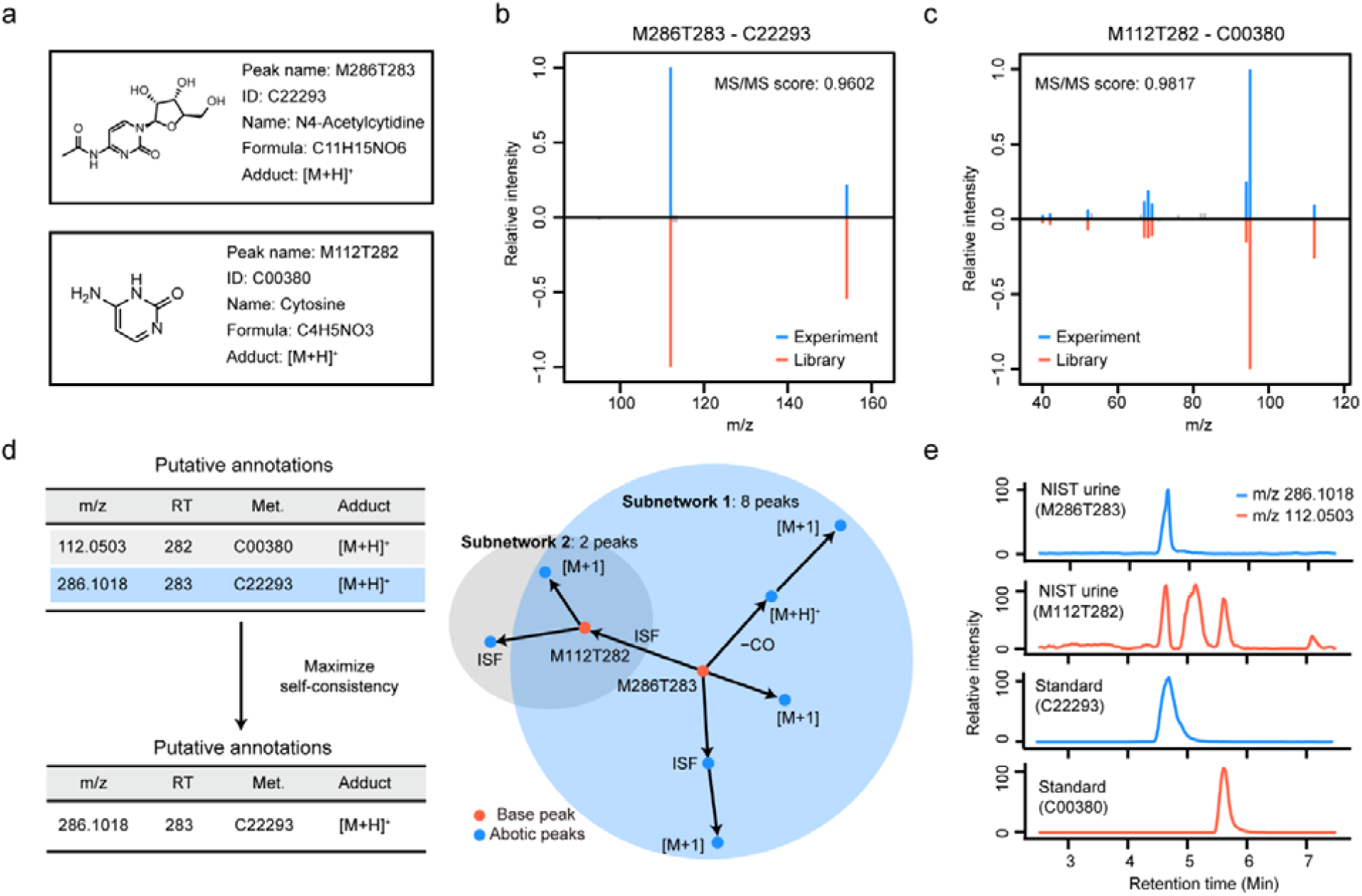
KGMN recognized the in-source fragments of N4-Acetylcytidine. (**a**) Peak M286T283 and peak M112T282 were annotated as N4-Acetylcytidine and cytosine in MetDNA1, respectively. (**b**-**c**) MS/MS spectral match between experimental MS/MS spectra and the standard spectral libraries for N4-Acetylcytidine (**b**) and cytosine (**c**). (**d**) Peak correlation subnetwork recognized M112T282 as an in-source fragment of M286T283. (**e**) The parallel acquisition of NIST human urine sample and chemical standards confirmed that peak M112T282 is an in-source fragment of M286T283.

**Extended Data Figure 4.**
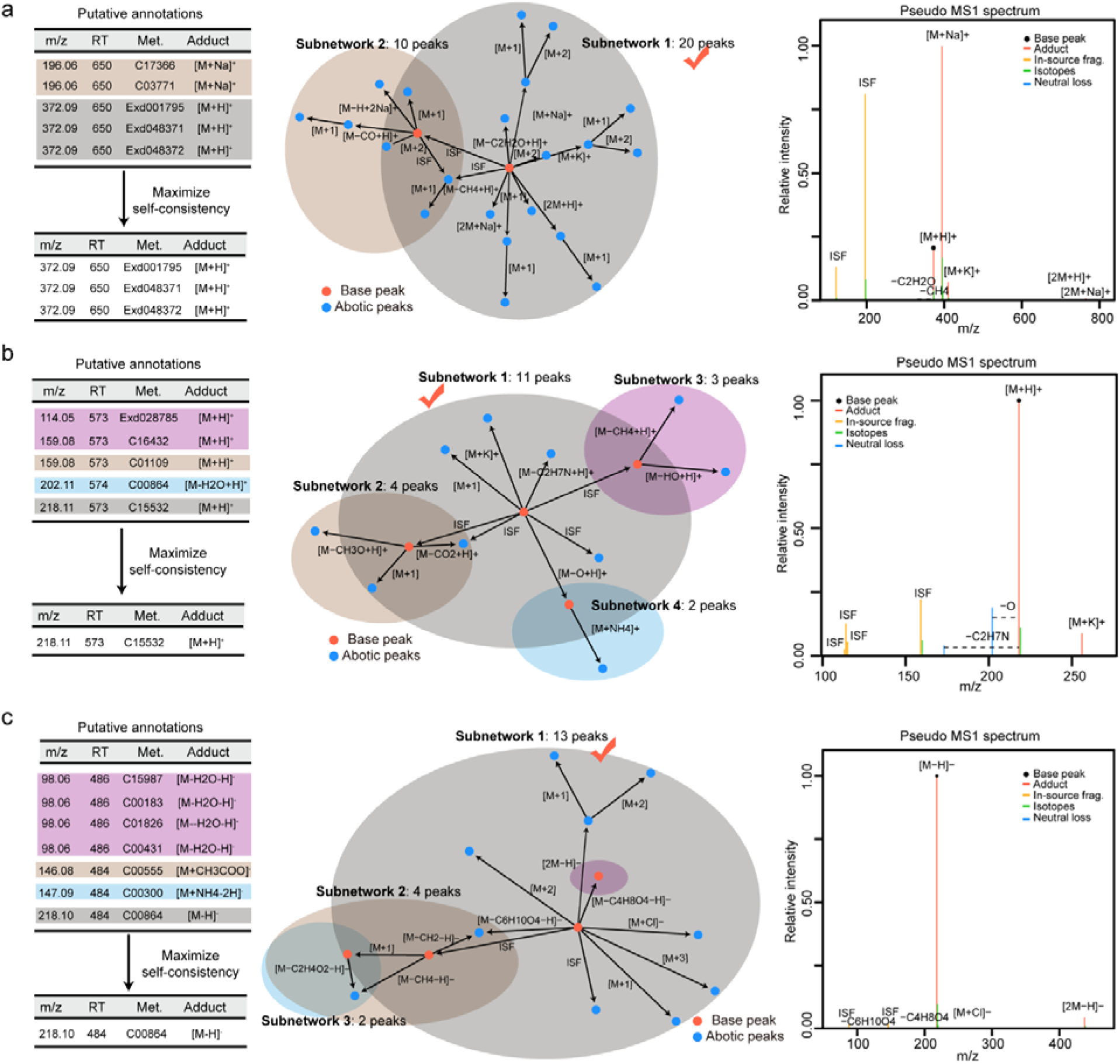
Examples of abiotic peak recognition and peak assignment in KGMN. (**a-c**) Abiotic peaks and putative annotations for (**a**) M372T650, (**b**) M218T573 and (**c**) M218T484. The left panel is the table for the reduction of putative annotations; the middle panel is the conflicted peak correlation subnetworks; the right panel is the pseudo MS1 spectrum after resolving the conflict peak correlations. The examples were retrieved from NIST human urine samples.

**Extended Data Figure 5.**
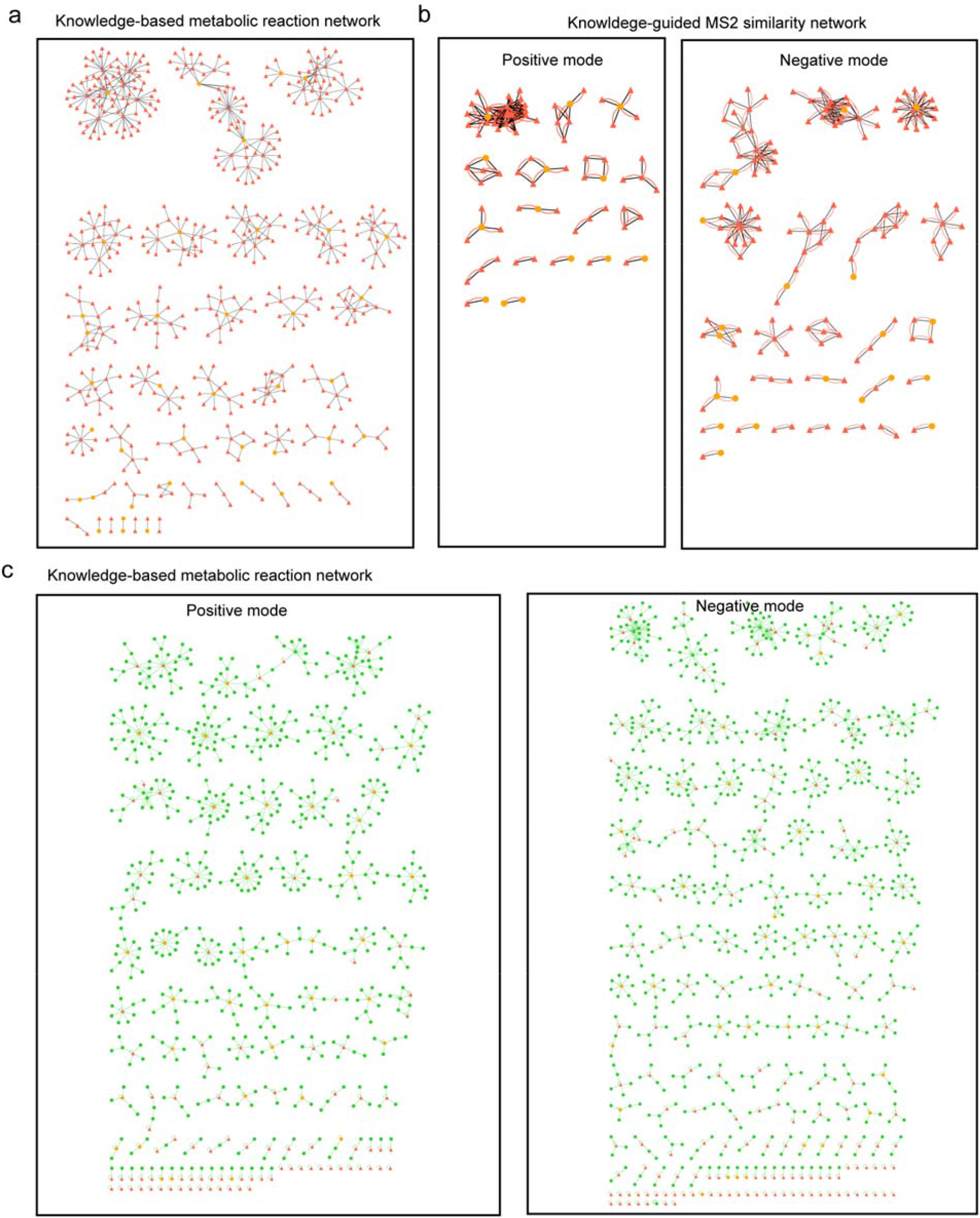
Knowledge-guided multi-layer networks of 46std_mix data sets. (**a**) Knowledge-based metabolic reaction network of 46 seed metabolites and unknown metabolites. The orange and red nodes represent seed and unknown metabolites, respectively. The unknown metabolites were curated via *in-silico* enzymatic reactions. The edges represent a biotransformation from known reactions or *in-silico* reactions. This network contains 531 unknown structures and 642 reaction pairs. (**b**) Knowledge-guided MS2 similarity network of 46 seed metabolites and unknown metabolites. The black and red edges represent the biotransformation and MS/MS spectral similarity. The edge of biotransformation represents two nodes can be transformed within 3-step reactions. The edge of MS/MS spectral similarity represents two nodes having MS/MS similarity (dot product score ≥0.5) or shared fragments (n≥4). Only linked peaks are showed here. (**c**) Global peak correlation network of 46std_mix data sets. The orange, red and green nodes represent seed, unknown and abiotic peaks. The edge represents an abiotic relationship (isotope, adduct, neutral loss or in-source fragment) between two nodes. A total of 745 and 932 peaks are included in positive and negative modes, respectively.

**Extended Data Figure 6.**
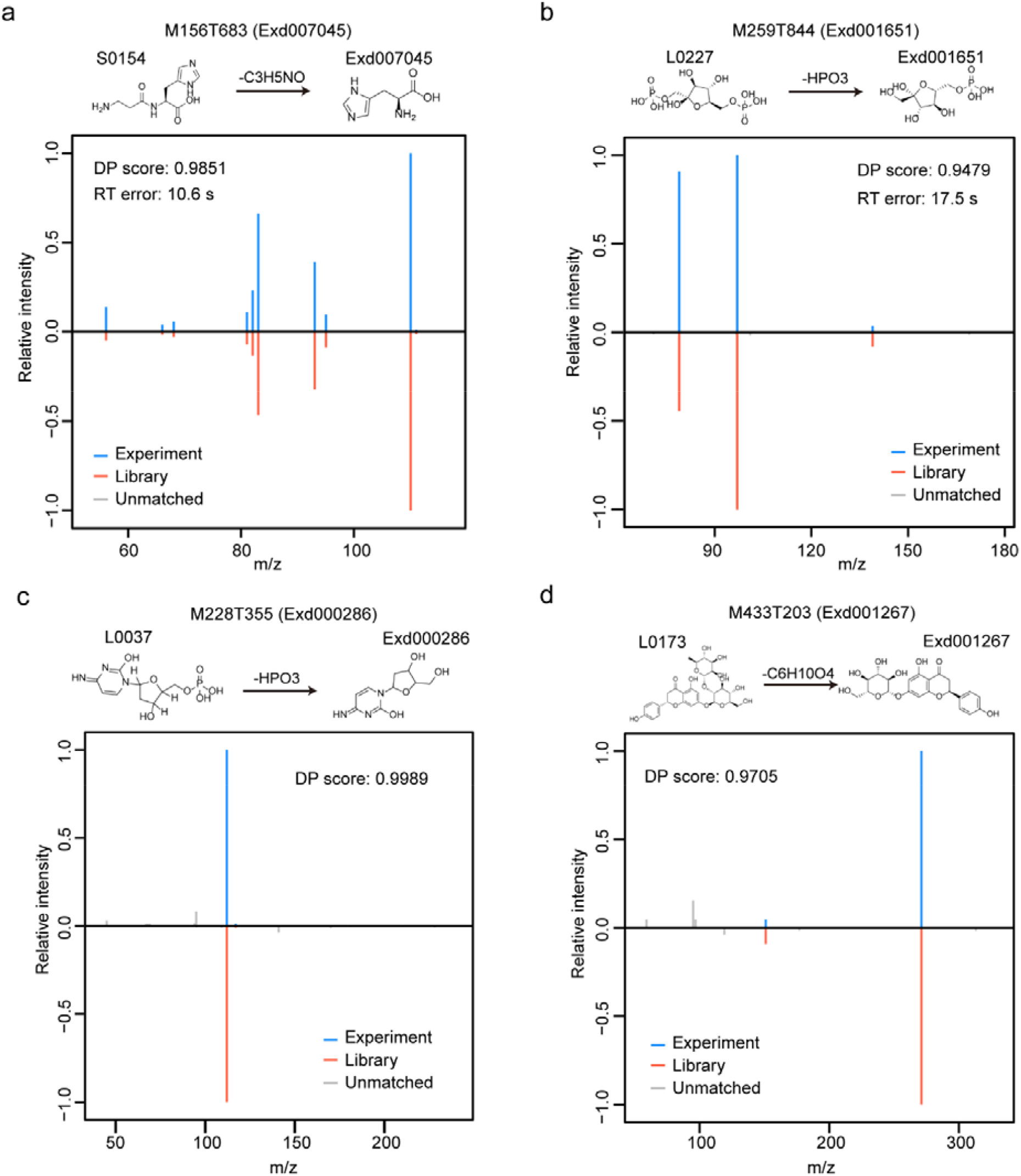
Validation examples of annotated unknowns in 46std_mix data sets. (**a**-**b**) Validation of unknowns using standards: (**a**) M156T683 (Exd007045, L-Histidine); and (**b**) M259T844 (Exd001651, D-Fructose 6-phosphate) in positive and negative modes, respectively; (**c**-**d**) validation of unknowns using public spectral databases: (**c**) 28T355 (Exd000286, Deoxycytidine), and (**d**) M433T203 (Exd001267, Naringenin 7-O-beta-D-glucoside) through MoNA and Metlin databases, respectively.

**Extended Data Figure 7.**
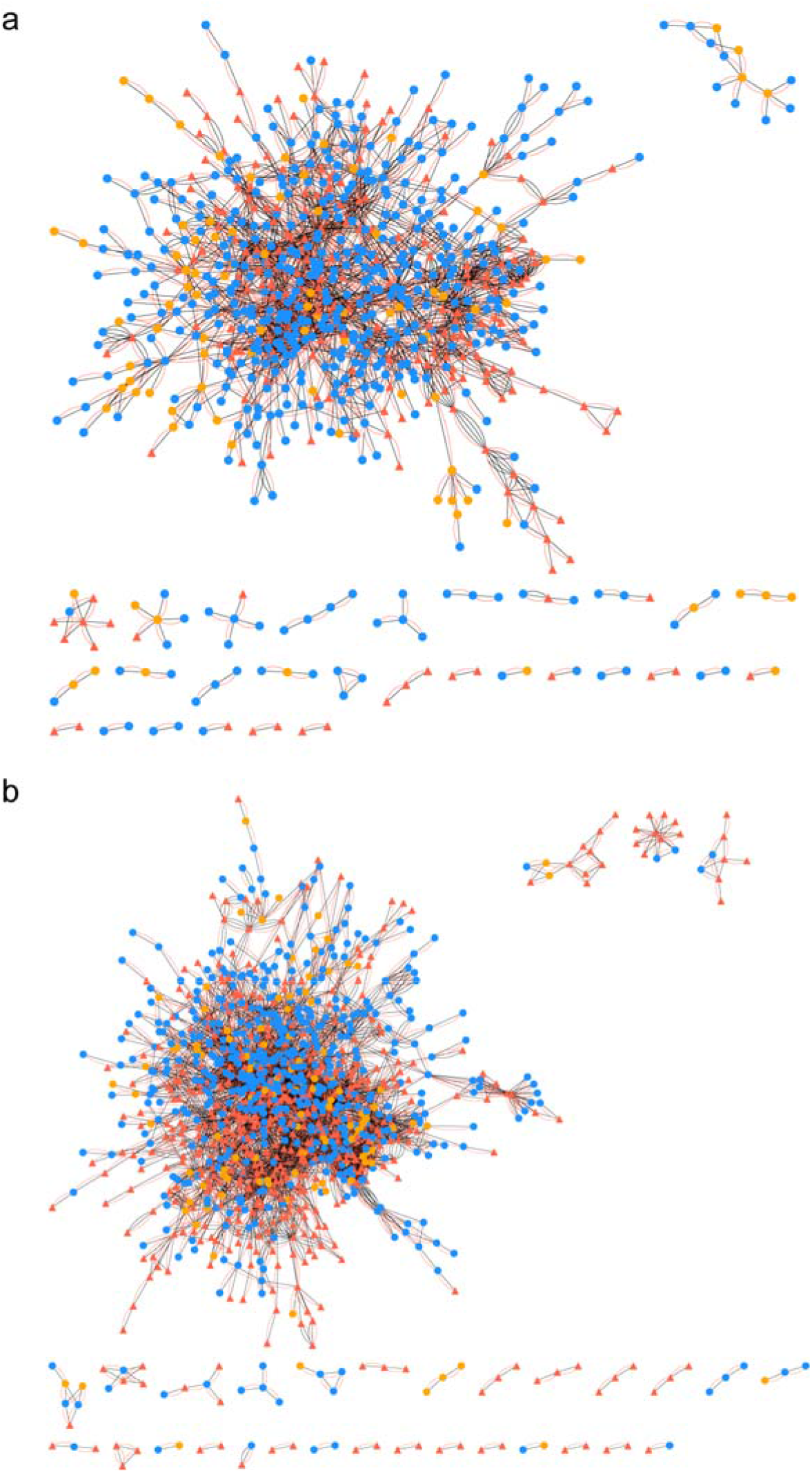
Knowledge-guided MS/MS similarity network of NIST human urine sample: (**a**) positive mode; (b) negative mode. The positive mode network contains 1,100 nodes, and 3,170 edges. The negative mode contains 1,444 nodes, and 7,810 edges. The orange, blue, and red nodes represent seed, known and unknown metabolites, respectively. The black and red edges represent the biotransformation edge and the MS/MS similarity edge, respectively. The edge of biotransformation represents two nodes can be transformed within 3-step reactions. The edge of MS/MS similarity represents two nodes having MS/MS similarity (dot product score ≥ 0.5) or shared fragments (n ≥ 4). Only linked peaks are showed here.

**Extended Data Figure 8.**
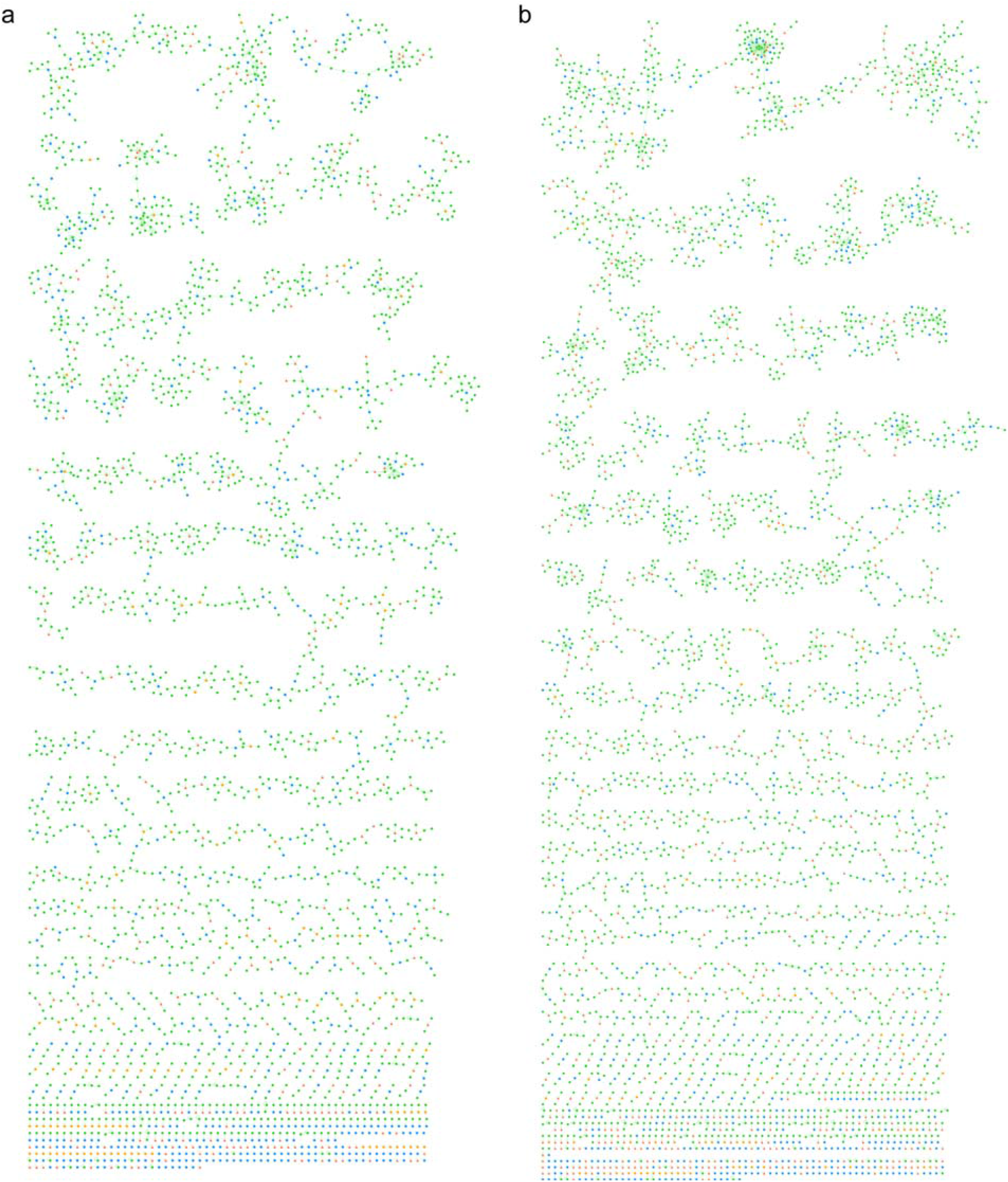
Global peak correlation network of NIST human urine sample in positive (**a**) and negative (**b**) modes. It contains 3,301 nodes and 4,374 edges in positive mode, and 4,117 nodes and 5,750 edges in negative mode. The orange, blue, and red nodes represent seed, known and unknown metabolites from network 2, which were used as base peaks here. The green nodes represent abiotic peaks.

**Extended Data Figure 9.**
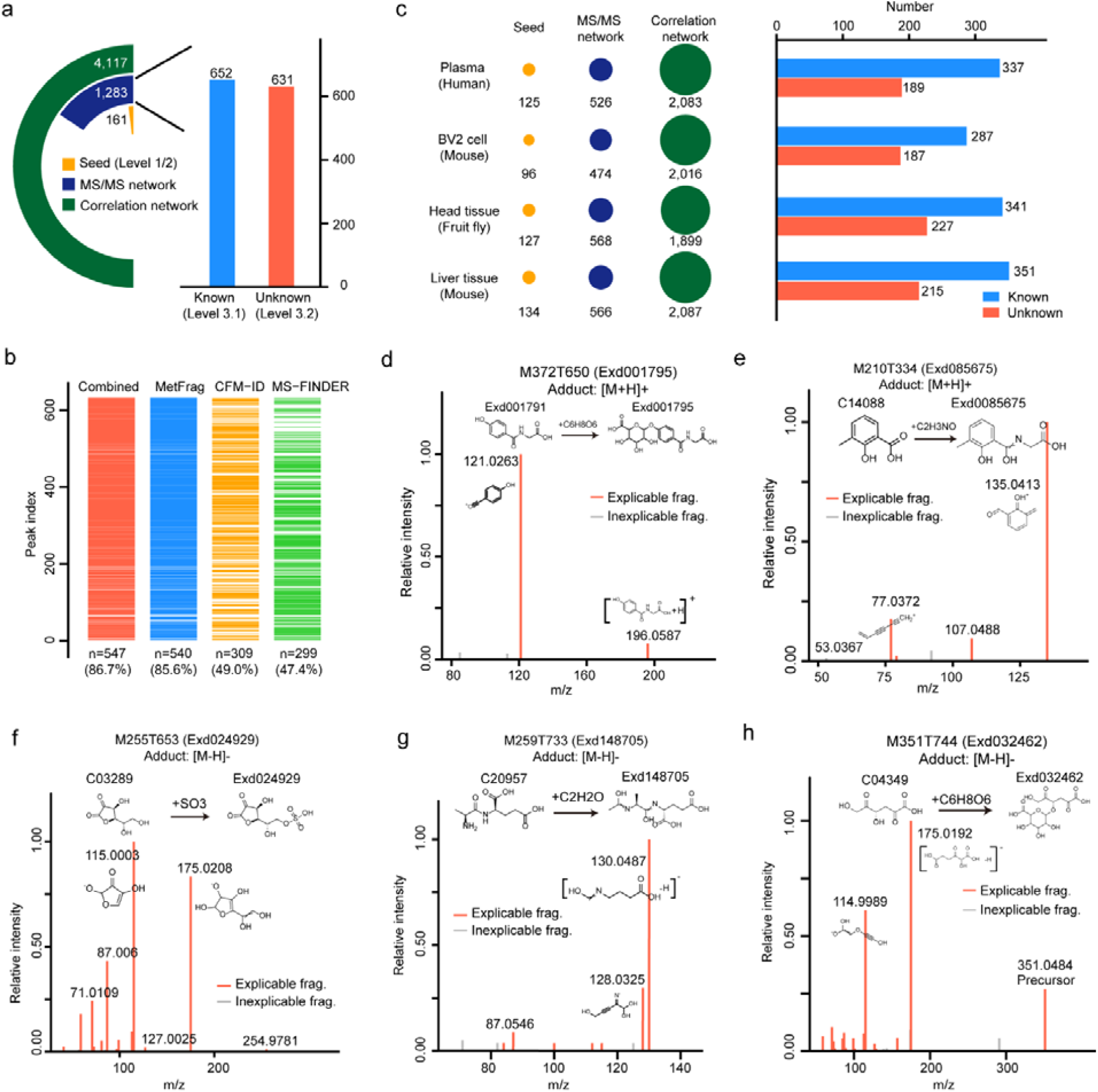
Global annotation of unknown metabolites in negative mode and validation examples of unknowns using *in-silico* MS/MS tools. (**a**) The annotated known and unknown metabolites in NIST human urine samples in negative mode. The left panel is the statistics of annotated peaks in the multi-layer networks, and the right panel is the statistics of annotated known and unknown peaks. (**b**) Validations of annotated unknown metabolites in urine samples with different *in-silico* MS/MS tools. (**c**) Global annotations of metabolites in different biological samples in negative mode. The left panel is the statistics of annotated peaks in the multi-layer network, and the right panel is the statistics of known and unknown metabolites. (**d**-**h**) Validation examples of unknown metabolites using *in-silico* MS/MS tools.

**Extended Data Figure 10.**
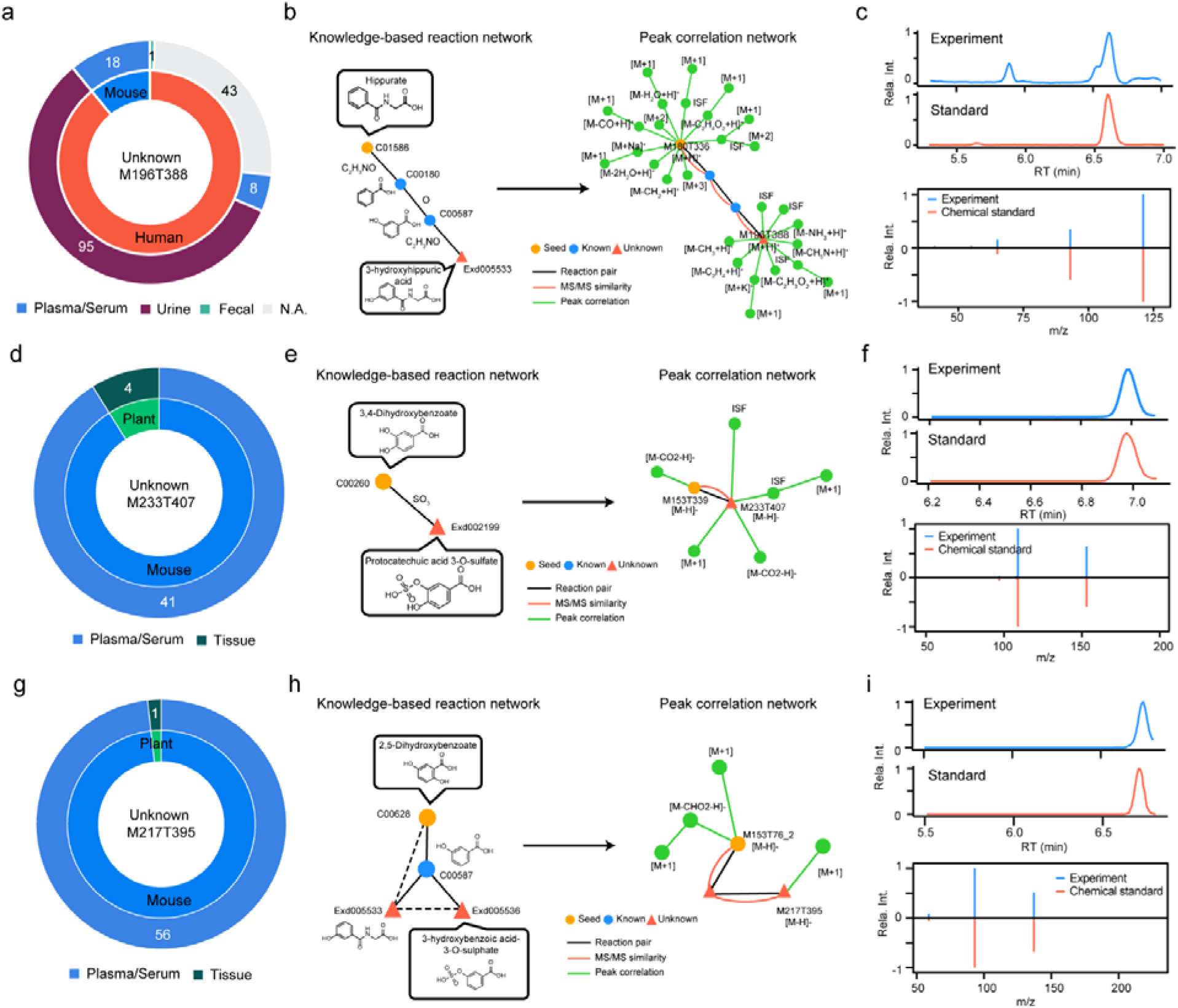
The repository-mining and structural validations of 3 recurrent unknown metabolites. (**a**-**c**) a recurrent unknown metabolite (M196T388, 3-hydroxyhippuric acid); (**d**-**g**) a recurrent unknown metabolite (M233T407, protocatechuic acid 3-O-sulfate); (**g**-**i**) a recurrent unknown metabolite (M217T395, 3-hydroxybenzoic acid-3-O-sulphate). The panels **a, d, g** are recurrent distributions of species and sample types; the inner and outer pie plots are the distributions in species and sample types, respectively. The panels **b, e, h** are the details of unknown annotations using KGMN. The panels **c, f, i** are the structural validations using the synthetic standards by matching the retention time and MS/MS spectra.

## References

1. Wishart, D. S. Emerging applications of metabolomics in drug discovery and precision medicine. Nat. Rev. Drug Discov. 15, 473–484 (2016).

2. Jang, C., Chen, L. & Rabinowitz, J. D. Metabolomics and Isotope Tracing. Cell 173, 822–837 (2018).

3. Bauermeister, A., Mannochio-Russo, H., Costa-Lotufo, L. V., Jarmusch, A. K. & Dorrestein, P. C. Mass spectrometry-based metabolomics in microbiome investigations. Nat. Rev. Microbiol. (2021) doi:10.1038/s41579-021-00621-9.

4. Giera, M., Yanes, O. & Siuzdak, G. Metabolite discovery: Biochemistry’s scientific driver. Cell Metab. 34, 21–34 (2022).

5. Tsugawa, H., Rai, A., Saito, K. & Nakabayashi, R. Metabolomics and complementary techniques to investigate the plant phytochemical cosmos. Nat. Prod. Rep. 38, 1729–1759 (2021).

6. Aurich, D., Miles, O. & Schymanski, E. L. Historical exposomics and high resolution mass spectrometry. Exposome 1, (2021).

7. Perez de Souza, L., Alseekh, S., Scossa, F. & Fernie, A. R. Ultra-high-performance liquid chromatography high-resolution mass spectrometry variants for metabolomics research. Nat. Methods 18, 733–746 (2021).

8. Alseekh, S. et al. Mass spectrometry-based metabolomics: a guide for annotation, quantification and best reporting practices. Nat. Methods 18, 747–756 (2021).

9. Domingo-Almenara, X., Montenegro-Burke, J. R., Benton, H. P. & Siuzdak, G. Annotation: A Computational Solution for Streamlining Metabolomics Analysis. Anal. Chem. 90, 480–489 (2018).

10. Sindelar, M. & Patti, G. J. Chemical Discovery in the Era of Metabolomics. J. Am. Chem. Soc. 142, 9097–9105 (2020).

11. Jarmusch, S. A., Van Der Hooft, J. J. J., Dorrestein, P. C. & Jarmusch, A. K. Advancements in capturing and mining mass spectrometry data are transforming natural products research. Nat. Prod. Rep. 38, 2066–2082 (2021).

12. Vinaixa, M. et al. Mass spectral databases for LC/MS-and GC/MS-based metabolomics: State of the field and future prospects. TrAC - Trends Anal. Chem. 78, 23–35 (2016).

13. Kind, T. et al. Identification of small molecules using accurate mass MS/MS search. Mass Spectrom. Rev. 37, 513–532 (2018).

14. Tsugawa, H. et al. A cheminformatics approach to characterize metabolomes in stable-isotope-labeled organisms. Nat. Methods 16, 295–298 (2019).

15. Ruttkies, C., Schymanski, E. L., Wolf, S., Hollender, J. & Neumann, S. MetFrag relaunched: incorporating strategies beyond in silico fragmentation. J. Cheminform. 8, 3 (2016).

16. Allen, F., Pon, A., Wilson, M., Greiner, R. & Wishart, D. CFM-ID: A web server for annotation, spectrum prediction and metabolite identification from tandem mass spectra. Nucleic Acids Res. 42, 94–99 (2014).

17. Tsugawa, H. et al. Hydrogen Rearrangement Rules: Computational MS/MS Fragmentation and Structure Elucidation Using MS-FINDER Software. Anal. Chem. 88, 7946–7958 (2016).

18. Dührkop, K. et al. SIRIUS 4: a rapid tool for turning tandem mass spectra into metabolite structure information. Nat. Methods 16, 299–302 (2019).

19. Wishart, D. S. et al. HMDB 4.0: The human metabolome database for 2018. Nucleic Acids Res. 46, D608–D617 (2018).

20. Kim, S. et al. PubChem 2019 update: Improved access to chemical data. Nucleic Acids Res. 47, D1102–D1109 (2019).

21. Hoffmann, M. A. et al. High-confidence structural annotation of metabolites absent from spectral libraries. Nat. Biotechnol. (2021) doi:10.1038/s41587-021-01045-9.

22. Wang, M. et al. Sharing and community curation of mass spectrometry data with Global Natural Products Social Molecular Networking. Nat. Biotechnol. 34, 828–837 (2016).

23. Shen, X. et al. Metabolic reaction network-based recursive metabolite annotation for untargeted metabolomics. Nat. Commun. 10, 1516 (2019).

24. Chen, L. et al. Metabolite discovery through global annotation of untargeted metabolomics data. Nat. Methods 18, 1377–1385 (2021).

25. da Silva, R. R. et al. Propagating annotations of molecular networks using in silico fragmentation. PLoS Comput. Biol. 14, 1–26 (2018).

26. van der Hooft, J. J. J., Wandy, J., Barrett, M. P., Burgess, K. E. V. & Rogers, S. Topic modeling for untargeted substructure exploration in metabolomics. Proc. Natl. Acad. Sci. 113, 13738–13743 (2016).

27. Ernst, M. et al. Molnetenhancer: Enhanced molecular networks by integrating metabolome mining and annotation tools. Metabolites 9, (2019).

28. Reily, M. D. et al. Proposed minimum reporting standards for chemical analysis. Metabolomics 3, 211–221 (2007).

29. Smith, C. A., Want, E. J., O’Maille, G., Abagyan, R. & Siuzdak, G. XCMS: Processing mass spectrometry data for metabolite profiling using nonlinear peak alignment, matching, and identification. Anal. Chem. 78, 779–787 (2006).

30. Tsugawa, H. et al. A lipidome atlas in MS-DIAL 4. Nat. Biotechnol. 38, 1159–1163 (2020).

31. Pluskal, T., Castillo, S., Villar-Briones, A. & Orešič, M. MZmine 2: Modular framework for processing, visualizing, and analyzing mass spectrometry-based molecular profile data. BMC Bioinformatics 11, 395 (2010).

32. Wang, M. et al. Mass spectrometry searches using MASST. Nat. Biotechnol. 38, 23–26 (2020).

33. Lai, Z. et al. Identifying metabolites by integrating metabolome databases with mass spectrometry cheminformatics. Nat. Methods 15, 53–56 (2018).

34. Huber, F. et al. Spec2Vec: Improved mass spectral similarity scoring through learning of structural relationships. PLoS Comput. Biol. 17, (2021).

35. Xing, S. et al. Retrieving and Utilizing Hypothetical Neutral Losses from Tandem Mass Spectra for Spectral Similarity Analysis and Unknown Metabolite Annotation. Anal. Chem. 92, 14476–14483 (2020).

36. Li, Y. et al. Spectral entropy outperforms MS/MS dot product similarity for small-molecule compound identification. Nat. Methods 18, 1524–1531 (2021).

37. Zhou, Z. et al. Ion mobility collision cross-section atlas for known and unknown metabolite annotation in untargeted metabolomics. Nat. Commun. 11, 1–13 (2020).

38. Hafner, J., Mohammadipeyhani, H., Sveshnikova, A., Scheidegger, A. & Hatzimanikatis, V. Updated ATLAS of Biochemistry with New Metabolites and Improved Enzyme Prediction Power. ACS Synth. Biol. 9, 1479–1482 (2020).

39. Tian, S. et al. CyProduct: A Software Tool for Accurately Predicting the Byproducts of Human Cytochrome P450 Metabolism. J. Chem. Inf. Model. 61, 3128–3140 (2021).

40. Jeffryes, J. G. et al. Chemical-damage MINE: A database of curated and predicted spontaneous metabolic reactions. Metab. Eng. 69, 302–312 (2022).

41. Yang, X., Neta, P. & Stein, S. E. Quality control for building libraries from electrospray ionization tandem mass spectra. Anal. Chem. 86, 6393–6400 (2014).

42. Li, H., Cai, Y., Guo, Y., Chen, F. & Zhu, Z. J. MetDIA: Targeted Metabolite Extraction of Multiplexed MS/MS Spectra Generated by Data-Independent Acquisition. Anal Chem 88, 8757–8764 (2016).

43. Kanehisa, M., Goto, S., Furumichi, M., Tanabe, M. & Hirakawa, M. KEGG for representation and analysis of molecular networks involving diseases and drugs. Nucleic Acids Res. 38, 355–360 (2009).

44. Djoumbou-Feunang, Y. et al. BioTransformer: A comprehensive computational tool for small molecule metabolism prediction and metabolite identification. J. Cheminform. 11, 1–25 (2019).

45. Liu, K. H. et al. Large scale enzyme based xenobiotic identification for exposomics. Nat. Commun. 12, 1–9 (2021).

46. Cai, Y., Weng, K., Guo, Y., Peng, J. & Zhu, Z.-J. An integrated targeted metabolomic platform for high-throughput metabolite profiling and automated data processing. Metabolomics 11, 1575–1586 (2015).

