## Supplementary Figures 1-3 and Tables 1-5 for "Metabolite annotation from knowns to unknowns through knowledge-guided multi-layer metabolic network"

**List for Supplementary data**

**Supplementary data 1**: Peak annotation evaluation between MetDNA1 and KGMN (MetDNA2)

**Supplementary data 2**: 46 standard mixture (46std_mix) and the knowledge-based metabolic reaction network

**Supplementary data 3**: KGMN results of 46std_mix data set and validation results

**Supplementary data 4**: KGMN results of NIST urine data sets and validation results

**Supplementary data 5**: KGMN results of different biological samples

**Supplementary data 6**: Recurrent unknowns of NIST urine via repository-mining

**Supplementary data 7**: Table of adducts, neutral losses, empirical rules in KGMN

**Supplementary Figures 1-3 and Tables 1-5**

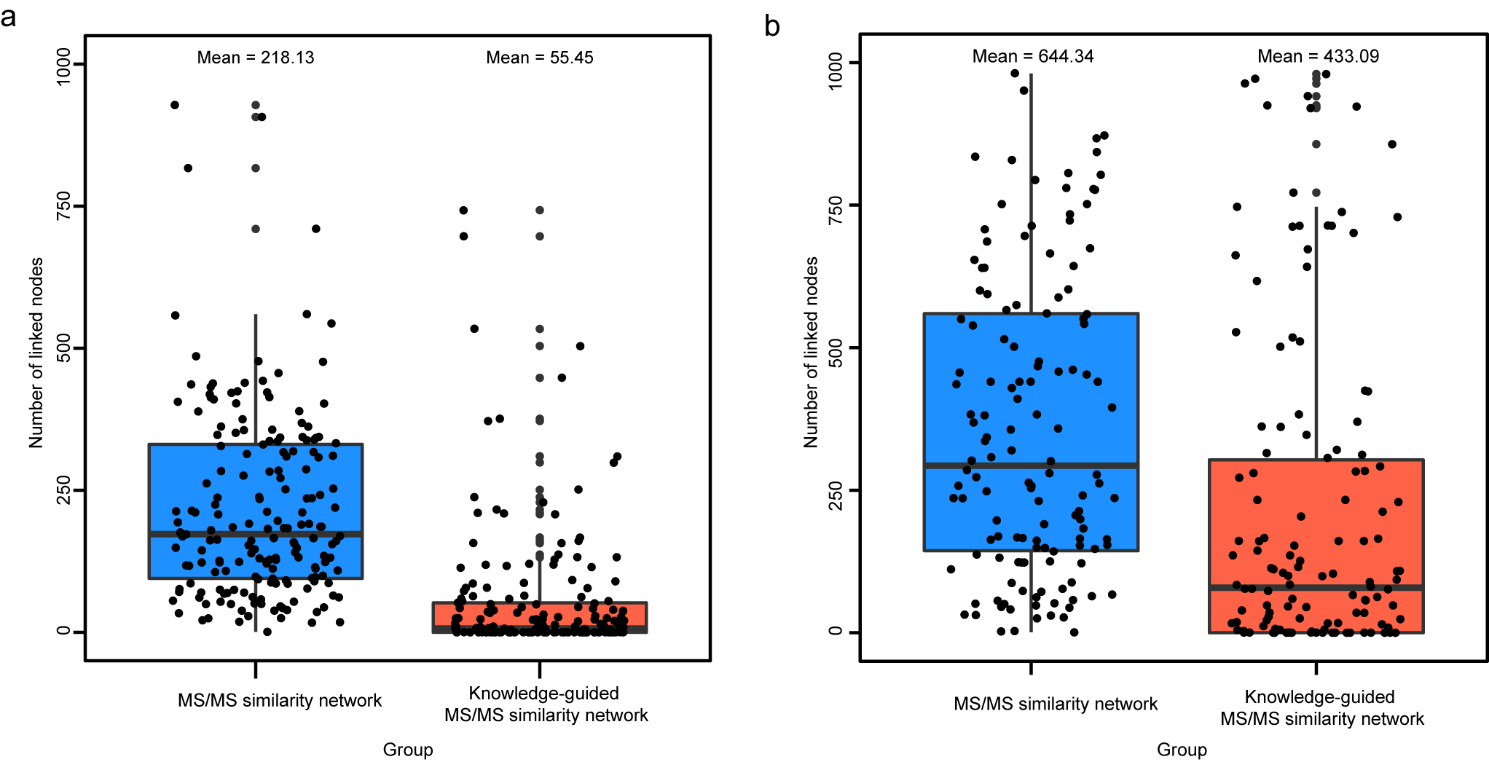

**Supplementary Figure 1.** Statistics of linked nodes in MS/MS similarity network or knowledge-guided MS/MS similarity network in positive (**a**) and negative modes (**b**), respectively. The linked nodes from seed metabolites in NIST human urine sample (N=181 and 163 in positive and negative modes, respectively) were included here. The cutoff of MS/MS similarity score is defined as 0.5. Neighbor metabolites within 3 steps were considered in knowledge-guided MS/MS similarity network.

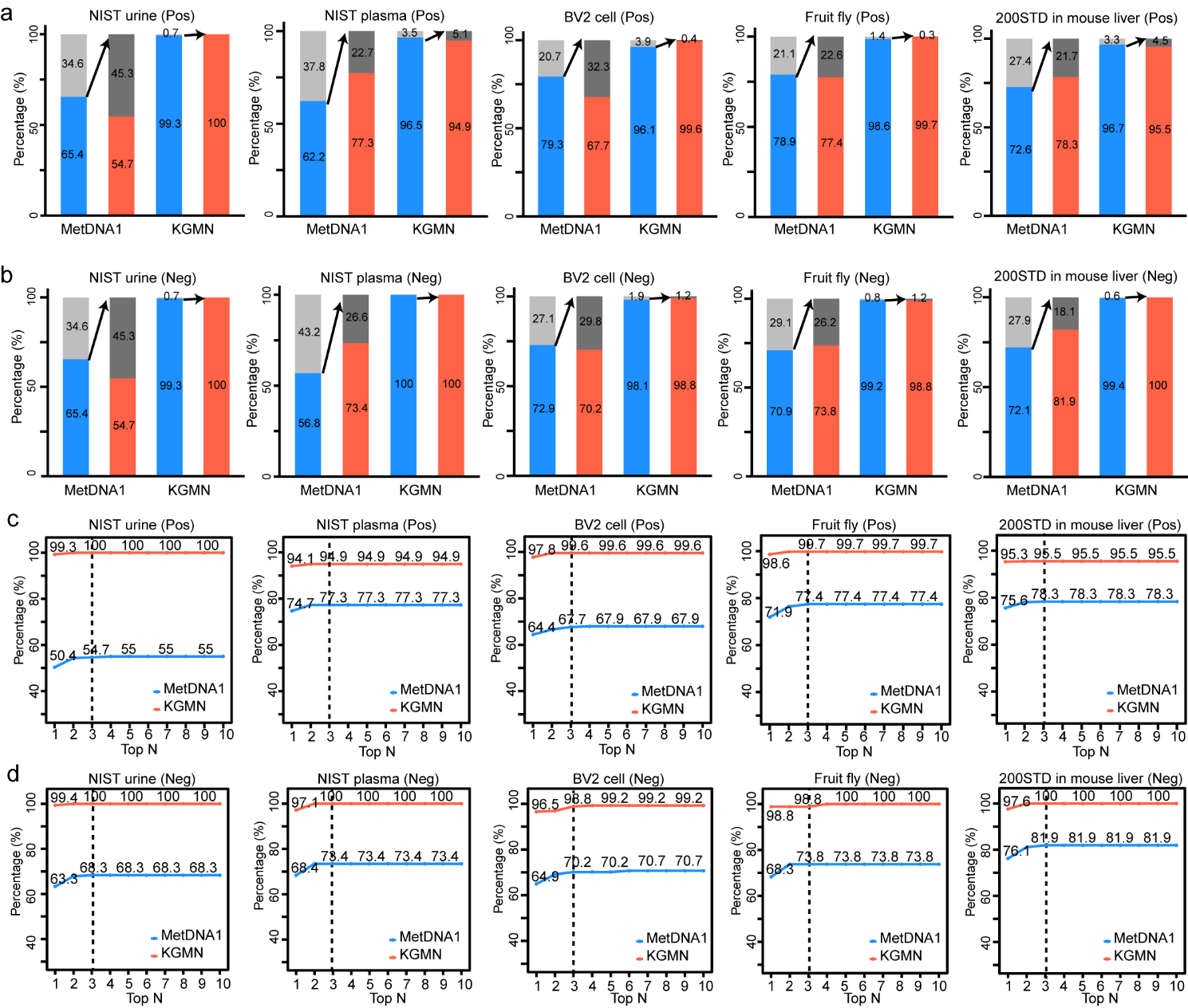

**Supplementary Figure 2.** Comparison between MetDNA1 and KGMN in different biological samples, including NIST human urine, NIST human plasma, BV2 cells, head tissues of fruit fly, and 200STD spiked mouse liver tissues. (**a**-**b**) Comparison of annotation coverages and correct/error percentages between MetDNA1 and KGMN in positive (**a**) and negative modes (**b**), respectively. (**c**-**d**) Correct and error rates among top n (n = 1 to 10) annotations in different biological samples in positive (**c**) and negative modes (**d**), respectively.

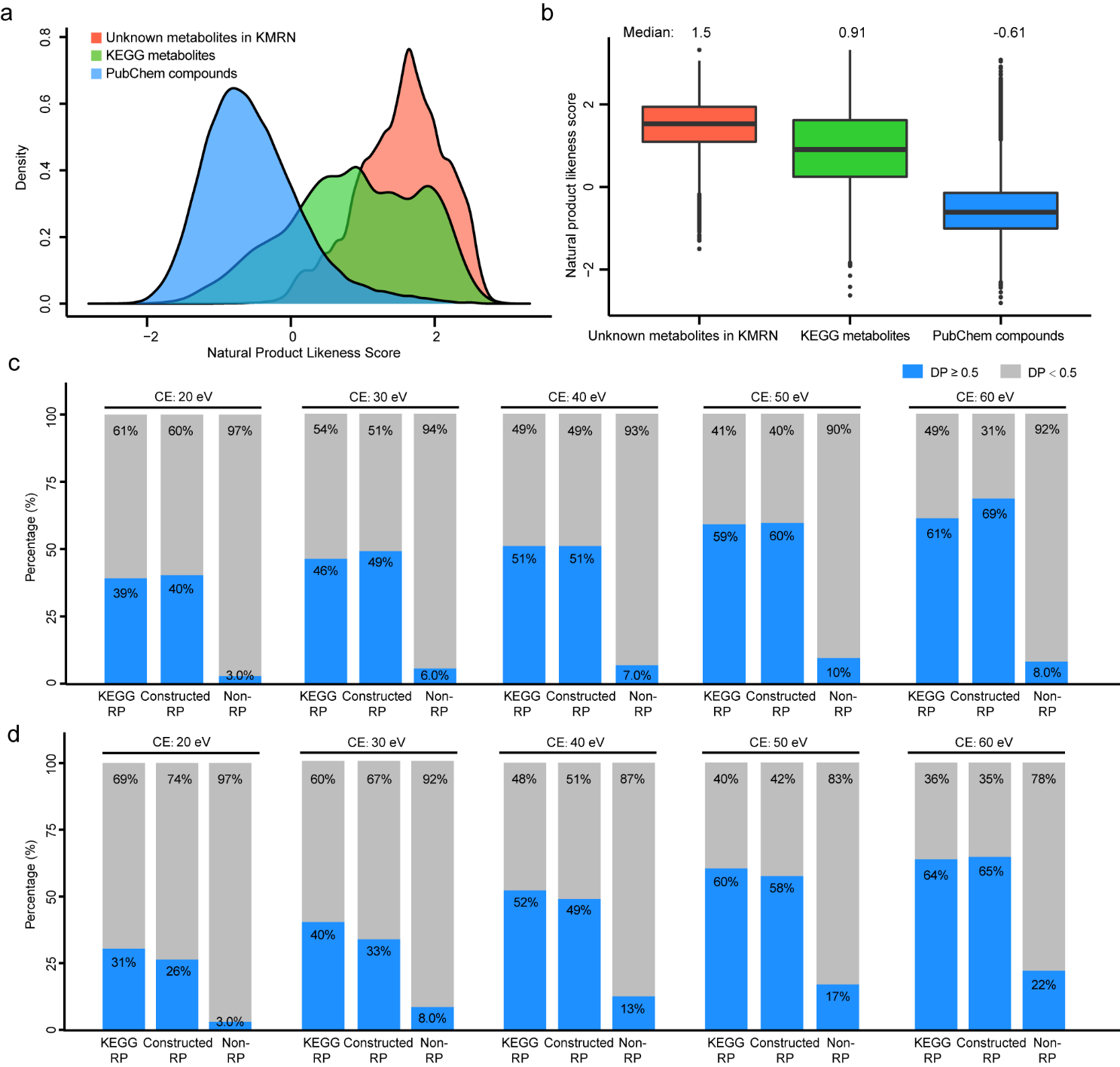

**Supplementary Figure 3.** Curated unknown metabolites and reaction pairs in the knowledge-based metabolic reaction network (KMRN). (**a**) Distribution of natural product likeness score of unknown metabolites in KMRN, KEGG metabolites, and PubChem compounds. 100,000 PubChem compounds were randomly retrieved to represent the PubChem database. (**b**) Natural product likeness score of unknown metabolites in KMRN, KEGG metabolites, and PubChem compounds. (**c**-**d**) MS/MS spectral similarity comparison among KEGG reaction pairs, *in-silico* curated unknown reaction pairs (i.e., constructed RP), and non-reaction pairs in positive (**c**) and negative (**d**), respectively.

**Supplementary Table 1.** Statistics of global peak annotation optimization to improve annotation accuracy.

| No. | Data set  (Polarity) | Peaks | MetDNA1 | | | MetDNA2 | | |
| --- | --- | --- | --- | --- | --- | --- | --- | --- |
|  |  |  | Peak with candi. | Candi. | Accuracy  (Top3) | Peak with candi. | Candi. | Accuracy  (Top 3) |
| 1 | NIST urine  (Pos) | 425 | 278 | 596 | 152  (54.7%) | 422 | 464 | 422  (100%) |
| 2 | NIST urine  (Neg) | 325 | 221 | 423 | 151  (68.3%) | 313 | 316 | 313  (100%) |
| 3 | NIST plasma (Pos) | 368 | 229 | 361 | 177  (77.3%) | 355 | 392 | 337  (94.9%) |
| 4 | NIST plasma (Neg) | 139 | 79 | 129 | 58  (73.4%) | 139 | 153 | 139  (100%) |
| 5 | BV2 cell  (Pos) | 464 | 368 | 604 | 249  (67.7%) | 446 | 457 | 444  (99.6%) |
| 6 | BV2 cell  (Neg) | 262 | 191 | 307 | 134  (70.2%) | 257 | 286 | 254  (98.8%) |
| 7 | Fruit fly head  (Pos) | 365 | 288 | 442 | 223  (77.4%) | 360 | 383 | 359  (99.7%) |
| 8 | Fruit fly head  (Neg) | 258 | 183 | 353 | 135  (73.8%) | 256 | 280 | 253  (98.8%) |
| 9 | 200STD in mouse liver  (Pos) | 508 | 369 | 459 | 289  (78.3%) | 491 | 506 | 469  (95.5%) |
| 10 | 200STD in mouse liver  (Neg) | 337 | 243 | 361 | 199  (81.9%) | 335 | 356 | 335  (100%) |
| Summary | | 3,451 | 2,449 | 4,035 | 1,767 | 3,374 | 3,593 | 3,325 |

**Supplementary Table 2.** Statistics of biotransformation types in 46std_mix data set.

| No. | Biotransformation | Positive mode | Negative mode |
| --- | --- | --- | --- |
| 1 | SO3 | 41 | 91 |
| 2 | C6H8O6 | 12 | 64 |
| 3 | HPO3 | 21 | 20 |
| 4 | O | 10 | 12 |
| 5 | H2 | 11 | 8 |
| 6 | H2O | 3 | 8 |
| 7 | C2H2O | 1 | 7 |
| 8 | C2H3NO | 0 | 3 |
| 9 | C4H4O3 | 1 | 2 |
| 10 | CH2 | 0 | 2 |
| 11 | CH3 | 0 | 2 |
| 12 | C10H10N4O3 | 0 | 1 |
| 13 | C3H5NO | 2 | 1 |
| 14 | C6H10O4 | 0 | 1 |
| 15 | C6H9O6 | 0 | 1 |

**Supplementary Table 3**. Statistics of annotated peaks in different biological samples

| Data sets | Seed peaks | MS/MS network | | | Peak correlation network |
| --- | --- | --- | --- | --- | --- |
|  |  | Known | Unknown | Sum |  |
| NIST urine (Pos) | 173 | 634 | 293 | 927 | 3,301 |
| NIST urine  (Neg) | 161 | 652 | 631 | 1,283 | 4,117 |
| NIST plasma  (Pos) | 135 | 310 | 73 | 383 | 1,774 |
| NIST plasma  (Neg) | 125 | 337 | 189 | 526 | 2,083 |
| BV2 cell  (Pos) | 188 | 398 | 183 | 581 | 2,827 |
| BV2 cell  (Neg) | 96 | 287 | 187 | 474 | 2,016 |
| Fruit fly brain  (Pos) | 187 | 265 | 122 | 387 | 1,883 |
| Fruit fly brain  (Neg) | 127 | 341 | 227 | 568 | 1,899 |
| Mouse liver  (Pos) | 209 | 270 | 107 | 377 | 2,464 |
| Mouse liver  (Neg) | 134 | 351 | 215 | 566 | 2,087 |
| Average | 154 | 385 | 223 | 607 | 2,445 |

**Supplementary Table 4.** Statistics of unknown biotransformation types in NIST urine data set

| No. | Biotransformation | Pos | Neg | No. | Biotransformation | Pos | Neg |
| --- | --- | --- | --- | --- | --- | --- | --- |
| 1 | SO3 | 322 | 1045 | 31 | H | 2 | 3 |
| 2 | C6H8O6 | 353 | 905 | 32 | C19H20N3O11P | 2 | 2 |
| 3 | H2 | 251 | 505 | 33 | C29H49N3O17P2 | 0 | 2 |
| 4 | O | 71 | 160 | 34 | C3H3O5P | 0 | 2 |
| 5 | HPO3 | 15 | 119 | 35 | C5H8NO3 | 0 | 2 |
| 6 | H2O | 100 | 108 | 36 | CO | 6 | 2 |
| 7 | C2H2O | 60 | 105 | 37 | C12H22N2O7 | 0 | 1 |
| 8 | C2H3NO | 57 | 83 | 38 | C14H26O | 0 | 1 |
| 9 | CH2 | 41 | 59 | 39 | C15H9O4 | 4 | 1 |
| 10 | isomer | 33 | 56 | 40 | C18H14N2O7 | 0 | 1 |
| 11 | CH3 | 10 | 38 | 41 | C2H4O | 0 | 1 |
| 12 | C7H12O6 | 20 | 34 | 42 | C30H25O12 | 6 | 1 |
| 13 | C6H9O6 | 3 | 29 | 43 | C30H48O2 | 1 | 1 |
| 14 | C6H10O5 | 33 | 22 | 44 | C3H2O | 0 | 1 |
| 15 | C6H11O5 | 17 | 20 | 45 | C61H100O11P2 | 0 | 1 |
| 16 | CO2 | 10 | 19 | 46 | C67H110O16P2 | 0 | 1 |
| 17 | C11H18O10 | 0 | 10 | 47 | C6H13N4O | 0 | 1 |
| 18 | C7H10O6 | 8 | 10 | 48 | C8H13NO | 0 | 1 |
| 19 | C2O3 | 0 | 6 | 49 | HO3S | 0 | 1 |
| 20 | C15H9O5 | 3 | 4 | 50 | -2O+H | 12 | 0 |
| 21 | C23H34N4O19P2 | 7 | 4 | 51 | C3H2O3 | 6 | 0 |
| 22 | C2H4 | 2 | 4 | 52 | C33H50O8 | 5 | 0 |
| 23 | C5H7NO3 | 0 | 4 | 53 | C27H40O2 | 3 | 0 |
| 24 | C10H15N3O6S | 2 | 3 | 54 | C6H10O4 | 2 | 0 |
| 25 | C12H16O10 | 3 | 3 | 55 | C7H4O4 | 1 | 0 |
| 26 | C12H20O10 | 5 | 3 | 56 | C7H5NO | 1 | 0 |
| 27 | C15H8O2 | 6 | 3 |  |  |  |  |
| 28 | C3H6NO | 1 | 3 |  |  |  |  |
| 29 | C8H12O7 | 4 | 3 |  |  |  |  |
| 30 | CH6N7O15P3S | 0 | 3 |  |  |  |  |

**Supplementary Table 5**. Compound search with history versions of HMDB

| Items | | Version of HMDB | | |
| --- | --- | --- | --- | --- |
|  |  | HMDB 3.6 | HMDB 4.0 | HMDB 5.0 |
| Download date | | 2022-04-06 | 2022-04-06 | 2022-04-06 |
| Release date | | 2017-09-10 | 2018-12-18 | 2021-11-02 |
| Compound number | | 74,435 | 113,983 | 217,776 |
| 1 | o-sulfotyrosine | No | No | Yes |
| 2 | 4-hydroxyhippuric acid | Yes | Yes | Yes |
| 3 | 3-hydroxyhippuric acid | Yes | Yes | Yes |
| 4 | protocatechuic acid 3-O-sulfate | No | Yes | Yes |
| 5 | 3-hydroxybenzoic acid-3-O-sulphate | Yes | Yes | Yes |

Note: in the manuscript, we used HMDB 4.0 for unknown checking.
